# Decoding Early Psychoses: Unraveling Stable Microstructural Features Associated with Psychopathology Across Independent Cohorts

**DOI:** 10.1101/2024.05.10.593636

**Authors:** Haley R. Wang, Zhen-Qi Liu, Hajer Nakua, Catherine E. Hegarty, Melanie Blair Thies, Pooja K. Patel, Charles H. Schleifer, Thomas P. Boeck, Rachel A. McKinney, Danielle Currin, Logan Leathem, Pamela DeRosse, Carrie E. Bearden, Bratislav Misic, Katherine H. Karlsgodt

## Abstract

**Background:** Early Psychosis patients (EP, within 3 years after psychosis onset) show significant variability, making outcome predictions challenging. Currently, little evidence exists for stable relationships between neural microstructural properties and symptom profiles across EP diagnoses, limiting the development of early interventions.

**Methods:** A data-driven approach, Partial Least Squares (PLS) correlation, was used across two independent datasets to examine multivariate relationships between white matter (WM) properties and symptomatology, to identify stable and generalizable signatures in EP. The primary cohort included EP patients from the Human Connectome Project-Early Psychosis (n=124). The replication cohort included EP patients from the Feinstein Institute for Medical Research (n=78). Both samples included individuals with schizophrenia, schizoaffective disorder, and psychotic mood disorders.

**Results:** In both cohorts, a significant latent component (LC) corresponded to a symptom profile combining negative symptoms, primarily diminished expression, with specific somatic symptoms. Both LCs captured comprehensive features of WM disruption, primarily a combination of subcortical and frontal association fibers. Strikingly, the PLS model trained on the primary cohort accurately predicted microstructural features and symptoms in the replication cohort. Findings were not driven by diagnosis, medication, or substance use.

**Conclusions:** This data-driven transdiagnostic approach revealed a stable and replicable neurobiological signature of microstructural WM alterations in EP, across diagnoses and datasets, showing a strong covariance of these alterations with a unique profile of negative and somatic symptoms. This finding suggests the clinical utility of applying data-driven approaches to reveal symptom domains that share neurobiological underpinnings.

## Introduction

Psychotic spectrum disorders, such as schizophrenia and psychotic mood disorders, have been associated with overlapping symptom presentations (1) and shared etiology. Despite this, research into early psychosis (EP; within 3 years of onset) has predominantly remained compartmentalized within diagnostic categories. This approach has raised questions about the potential for uncovering shared neurobiological features (2,3), highlighting the need for methodologies that can robustly test findings across different populations. Further, a critical aspect of advancing our understanding in this field is the rigorous cross-validation of research findings across independent datasets. Such validation is essential for identifying robust biomarkers in neuroimaging, given the complexity and variability of psychotic disorders.

One hallmark of structural brain changes in psychosis is alterations in white matter (WM) (3–10), discernable with diffusion-weighted imaging (DWI). WM alterations, characterized by higher diffusivity and lower directional coherence, are consistent with disruptions in biological processes, such as neuro-inflammation and axonal density (11,12). These deficits are particularly pertinent to EP, as WM development in adolescence underlies the formation of higher-order cognitive functions vital for adaptive behavior, and could be key to the early disease process and functional outcomes in psychotic illness (13).

WM alterations in EP have been extensively investigated using DWI meta- and mega-analyses, commonly modeled by a diffusion tensor (DTI). These analyses have demonstrated widespread reductions of fractional anisotropy (FA; a putative index of WM coherence that has been associated with microstructural features such as myelination and fiber density) in individuals with psychosis compared to controls, with the largest effect sizes seen in corpus callosum, anterior corona radiata, and superior longitudinal fasciculus (SLF) (5,6,8). However, questions remain about the association of WM microstructure with various dimensions of psychopathology. While studies reported associations between FA reduction and positive, negative, and cognitive symptom severity (9), heterogeneity in both the symptoms and the regional distribution of FA reduction reported across studies have limited the conclusions to date. Heterogeneity between studies may be attributed to the reliance on univariate statistics, broadly summarized symptom domains, and relatively small samples (n<100) (14). Recently, these limitations have led the field towards multivariate and data-driven approaches for a more replicable and comprehensive understanding (15–17). Further, with symptom overlap across disorders, it is crucial to investigate brain-symptom relationships transdiagnostically, especially for adolescence and EP. EP is associated with diagnostic instability and highly variable prognosis, making categorical diagnoses less useful in this population and limiting clinical utility of traditional research approaches (1,18). Given these complexities, identifying shared neurobiological underpinnings of symptom profiles, agnostic to diagnosis (19), is pivotal for understanding the core neuropathological mechanism of psychotic disorders and to guide development of targeted treatments of unique symptom clusters for youth.

Our investigation employs Partial Least Squares (PLS) correlation, a statistical-learning method identifying optimally covarying patterns between high-dimensional features across modalities (17,19,20). This approach offers three distinct advantages. First, it reveals latent components (LCs) representing key covarying patterns between brain and clinical variables. Unlike univariate methods that leverage specific regions, PLS facilitates a nuanced, feature-level understanding of the interplay of how various brain metrics may be related to symptoms. Second, PLS efficiently manages collinearity, vital for delineating complex brain-symptom relationships (21,22). Third, PLS outputs facilitate flexible post-hoc analyses to systematically examine similarities and differences in brain-symptom correlations across diagnostic groups (23). Essentially, this methodology shifts focus from a single diagnostic univariate approach using *a priori* symptom measures to a transdiagnostic approach which allows subtyping clinical symptoms beyond traditional categories based on shared neurobiological features. This shift enables a more organic and granular investigation of psychosis neurobiology that capitalizes on the clinical heterogeneity of EP patient sample.

Here, we aimed to examine the multivariate relationship between WM microstructure alterations and psychopathology dimensions in two transdiagnostic EP samples. Specifically, we assessed whether the PLS-identified WM signature correlates with distinct symptom domains across EP diagnostic boundaries. We also established the signature’s stability, generalizability, and clinical utility through both external validation and prediction analysis using two independent EP datasets. To ensure its independence from potential confounders, we evaluated its links with factors such as antipsychotic medication load, substance use, and illness duration.

## Methods and Materials

### Sample characteristics

This study employed two independent EP datasets. The primary cohort, Human Connectome Project Early Psychosis Release 1.1 (HCP-EP), includes 124 patients aged 16-35 years (38.7% female). The replication cohort from the Multimodal Evaluation of Neural Disorders (MEND) Project (MH101506) includes 78 patients aged 16-25 years (24.4% female). Both cohorts included patients with diagnoses of schizophrenia, schizoaffective disorders, and psychotic mood disorders, within three years of onset. Complete demographic and clinical characteristics are summarized and compared in Table 1. See Supplemental Methods S1-4 in supplement for recruitment details. Diagnostic group differences in clinical scores are shown in Figure S1.

**Table 1.** Demographic, clinical, and medication characteristics of the primary and replication cohorts.

| Characteristics | HCP-EP<br>(primary analysis) | MEND<br>(replication analysis) | *P* Value |
| --- | --- | --- | --- |
| No. of participants | 124 | 78 | NA |
| Age, mean (SD), y | 22.7 (3.6) | 20.9 (2.5) | .01 |
| Male, No. (%) | 76 (61.3) | 59 (75.6) | .11 |
| Race, No. (%) |  |  |  |
| Asian | 8 (6.5) | 10 (12.8) | .15 |
| African American | 49 (39.5) | 32 (41) |  |
| American Indian/Alaska Native | 1 (0.8) | 1 (1.3) |  |
| White | 62 (50) | 24 (30.8) |  |
| Unknown or more than one race | 4 (3.2) | 11 (14.1) |  |
| Hispanic ethnicity, No. (%) | 8 (6.5) | 13 (16.7) | .08 |
| Parental education, mean (SD), y | 10.2 (2.3) | 10.8 (1.7) | .15 |
| WASI Full Scale IQ, mean (SD) | 101 (17.2) | 99.5 (14.7)<sup>a</sup> | .49 |
| Duration of Illness, mean (SD), y | 2.19 (1.4) | 2.1 (1.9)<sup>b</sup> | .52 |
| SCID diagnosis, No. (%) |  |  |  |
| Schizophrenia | 81 (65.3) | 39 (50) | .06 |
| Schizoaffective Disorder | 14 (11.3) | 23 (29.5) |  |
| Bipolar I Disorder or Major Depressive Disorder with psychotic features | 26 (21) | 16 (20.5) |  |
| Medication Use |  |  |  |
| Current dosage of chlorpromazine equivalent, mean (SD), mg | 159 (224.5) | 243.7 (192.2)<sup>c</sup> | .05 |
| Cannabis use in past 30 days, No. (%) | 16 (12.9) | 10 (15.9)<sup>d</sup> | .66 |
| YMRS score, mean (SD) | 4.9 (5.3) | 6.6 (7.3) | .10 |
| PANSS scores, mean (SD) |  |  |  |
| Positive | 11.2 (4) | NA | NA |
| Negative | 14 (5.4) |  |  |
| General | 24.7 (5.2) |  |  |
| BPRS score, mean (SD) |  |  |  |
| Total | NA | 31.2 (9.4) | NA |
| Total positive<sup>e</sup> |  | 13.9 (5.7) |  |
| SANS score, mean (SD) | NA | 18.6 (11.7) | NA |

*Note: WASI = Wechsler Abbreviated Scale of Intelligence; YMRS = Young Mania Rating Scale; PANSS = Positive and Negative Syndrome Scale; BPRS = Brief Psychiatric Rating Scale; SANS = Scale for the Assessment of Negative Symptoms; SCID = Structured Clinical Interview for DSM-IV.*
*NA indicates that the clinical scales were not used for the cohort.*
*Due to rounding percentages might not sum to 100.*
*Group characteristics were tested using two-sample t-tests for continuous and Chi-Square or Fisher's exact tests, as appropriate, for categorical variables.*
*a, b, c, d Observations missing covariate of interest = 10, 33, 17, 10, respectively.*
*<sup>e</sup> Total positive symptoms in the BPRS were calculated using items in the dimensions of positive and hostile-suspiciousness(80) (Specific symptoms are shown in Figure 3D and Table S1 in Supplement).*

### Microstructural features

Diffusion MRI data were preprocessed according to standard pipelines (Supplemental Methods S5-6). Mean values of 4 DTI metrics (FA, mean diffusivity (MD), axial diffusivity (AD), and radial diffusivity (RD)) were extracted for 48 WM tracts defined by the JHU White-Matter atlas (JHU-ICBM-DTI-81). In both HCP-EP and MEND datasets, 192 brain features (4 diffusion metrics across 48 tracts) were generated to capture a comprehensive microstructural profile of WM. Additionally, neuroCombat (24) was employed to harmonize multi-site diffusion data for HCP-EP to adjust for scanner differences at the feature level using the tract-averaged metrics.”

### Psychopathology features

We assessed 4 psychopathology dimensions: positive symptoms (e.g., hallucinations, delusions), negative symptoms (e.g., avolition, anhedonia), general psychopathology symptoms (e.g., tension, somatic concern) and mania symptoms (e.g., elevated mood, pressured speech). In HCP-EP, the Positive and Negative Syndrome Scale (PANSS) (25) and the Young Mania Rating Scale (YMRS) (26) were used, yielding 41 symptom-level features. In MEND, these same dimensions were assessed by the Brief Psychiatric Rating Scale (BPRS) (27), the Scale for the Assessment of Negative Symptoms (SANS) (28), and YMRS, resulting in 49 features (Table S2A-B).

### PLS analysis

We conducted independent PLS analyses for each dataset, deriving LCs that optimize the covarying patterns between brain and clinical features (Table S1-2). The PLS approach is particularly adept at managing multicollinearity, as it amplifies collections of variables that characterize the cross-product matrix, such as the covariance between DTI metrics like MD, RD, AD, and FA. This ensures that model fitting is not confounded by multicollinearity among these variables. While derived from the same eigenvalues of the diffusion tensor, these DTI metrics offer distinct insights into WM properties and their clinical implications. Before proceeding with the PLS models, we regressed out the effects of sex, age, and quadratic age using Ordinary Least Squares regression, using the residuals as input data. The subsequent PLS analyses decomposed the two data matrices into linearly independent sets of LCs characterized by maximum covariance. The resulting LCs, each composed of optimal weighing of original variables (e.g., raw symptom scores), were characterized by a corresponding singular value signifying the covariance explained between overall WM integrity and clinical symptoms (Supplemental Methods S7). By strategically handling within-block correlations and focusing on optimizing for covariance, PLS effectively manages the potential confounding effects of multicollinearity among DTI indexes.

As described in our previous work (29), significant LCs were determined using a permutation test with 5,000 iterations with false discovery rate (FDR) correction (q<.05). Projecting each patient’s raw data onto the LC yielded composite scores for both microstructural and clinical features encapsulated by that particular LC (analogous to factor scores). We then computed loadings as the Pearson’s correlation coefficient between the composite scores and each microstructural and clinical feature. These loadings indicate the shared variance between the original variables and the PLS score pattern, thereby reflecting the contribution level of each variable to the LCs. Confidence intervals for each loading were determined through cross-validation and bootstrapping (10,000 repetitions for microstructure and psychopathology LCs separately). Sensitivity analyses were implemented (Supplemental Methods S8); post-hoc analyses examined the impact of age, sex, IQ, sites, head motion, diagnoses, duration of illness, substance use, and medication on LC composite scores using Pearson’s correlation coefficient and two-sample t-tests (Supplemental Methods S9).

### Cross-cohort prediction

To test similarity of the LCs between cohorts, we used a cross-cohort prediction approach. Initially, we extracted the weights of each microstructural feature in the LCs from the PLS model trained by the primary cohort (HCP-EP). We then multiplied these PLS-derived weights from HCP-EP by the raw, individual-level microstructural metrics in the validation cohort (MEND) to compute the anticipated microstructural composite scores for each MEND patient. Subsequently, we assessed the correlation between the predicted microstructural composite scores of MEND and the empirical microstructural composite scores observed in MEND. Additionally, we examined the correlation between the predicted microstructural composite scores and the empirical clinical composite scores in MEND using Pearson’s correlation coefficient.

## Results

### PLS primary analysis in EP

In HCP-EP, PLS analysis revealed a single significant LC (LC_HCP_, p<.001) that captures 38.6% of the covariance between WM structures and clinical symptoms (Figure 1A, Figure S2). No post-FDR difference in microstructural composite scores was observed across diagnostic groups (q>.05; Figure 1B). However, individuals with schizophrenia spectrum disorders without mood features showed higher clinical composite scores (i.e., symptom severity) than schizoaffective (p<.05) and psychotic mood disorder groups (p<.01; Figure 1C). Top microstructural loadings were retained as features that contributed the most to LC_HCP_ (Figure 1D). Microstructural WM features with strongest loadings included higher values of MD and AD in a range of projection, commissural, brainstem and association tracts across the brain, representing a combination of tracts previously found in SZ studies such as the SLF and corona radiata, as well as other tracts such as the external and internal capsules and cingulum (Figure 1F; Table S3). Original FA features were negatively loaded, while MD, AD, and RD features were positively loaded onto the microstructural pattern. The difference in the direction of these relationships is expected, as higher FA typically reflects greater myelination or tract density, while increased MD, AD and RD are generally thought to reflect the opposite, signifying WM disruption. In clinical analyses, symptoms with significant loadings to the psychopathological pattern of LC_HCP_ are mainly negative symptoms, characterized by diminished emotional expression (Figure 1E). Tension positively loaded onto this psychopathological pattern, while decreased need for sleep—a symptom of mania—loaded negatively. This is consistent with the expressive deficits and hypoarousal often observed in negative symptoms. Overall, LC_HCP_ establishes a relationship between features of WM disruption and specific negative symptoms in EP.

**Figure 1.**
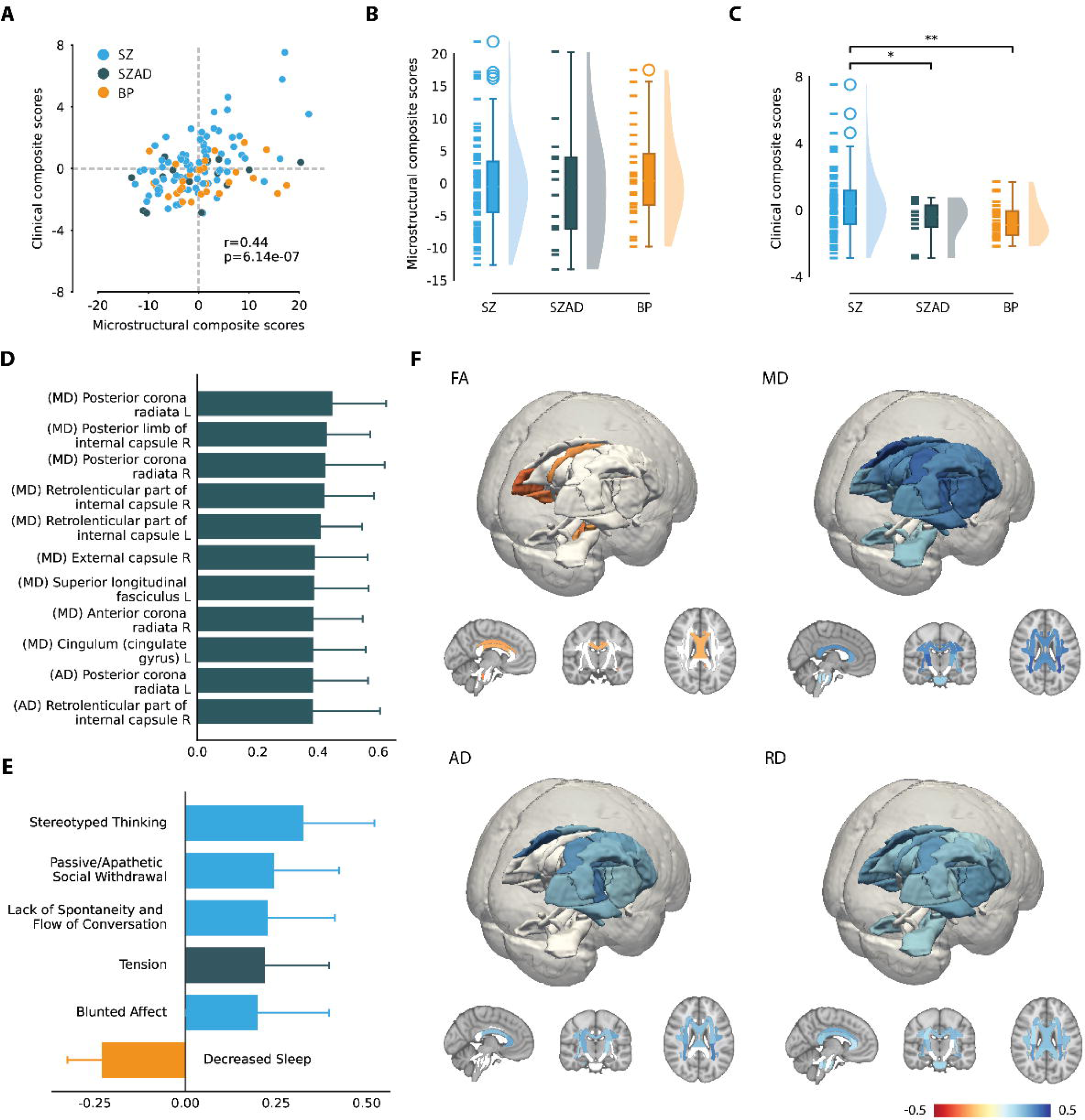
Primary Latent Component (LC) in HCP-EP reflects negative symptom dimension. **(A)** Correlation between individual-specific microstructural and clinical composite scores of patients. Scatterplot was color-coded by each primary diagnostic group. **(B)** Group differences in microstructural composite scores. **(C)** Group differences in clinical composite scores. Asterisks indicate t-tests that survived false discovery rate (FDR) correction (*=q<.05; **=q<.01). **(D)** Thresholded significant microstructural loadings of the LC. Strong loadings with an effect size greater than 2 standard deviations were shown in the bar chart. Error bars indicate bootstrapped standard deviations. **(E)** Unthresholded significant clinical loadings of the LC. Blue, green, and orange colors indicate the symptom dimensions of negative, general, and mania symptoms, respectively. **(F)** The loading coefficient of all significant microstructural features associated with the LC are projected to their corresponding WM tract location in the brain, categorized by DTI metrics. Non-significant tracts are in white.

### PLS replication analysis in MEND

Findings in MEND were consistent with HCP-EP findings, yielding a single significant LC (LC_MEND_, p=.01) that captured 37.8% of the covariance and showed a similar microstructural signature combining association tracts like the SLF with subcortical and brainstem tracts (Figure 2A; Figure S2). However, in MEND, FDR-corrected composite scores did not differ across diagnoses (q>.05; Figure 2B-C). The predominant microstructural features in LC_MEND_ showed increases in WM disruption, specifically elevated MD and RD in multiple tracts (Figure 2D; Figure 2F; Table S3). Clinically, LC_MEND_ positively correlated with loadings of diminished affective expressivity, characterized by fewer expressive gestures and less spontaneous movement (Figure 2E). Unique to this cohort was the positive loading of motor retardation with the psychopathological pattern, while symptoms of mania, such as increased sexual interest and elevated mood, were negatively loaded. This replication confirms the primary findings, linking features of WM disruption to select symptoms featuring hypoemotionality and hypoarousal in EP.

**Figure 2.**
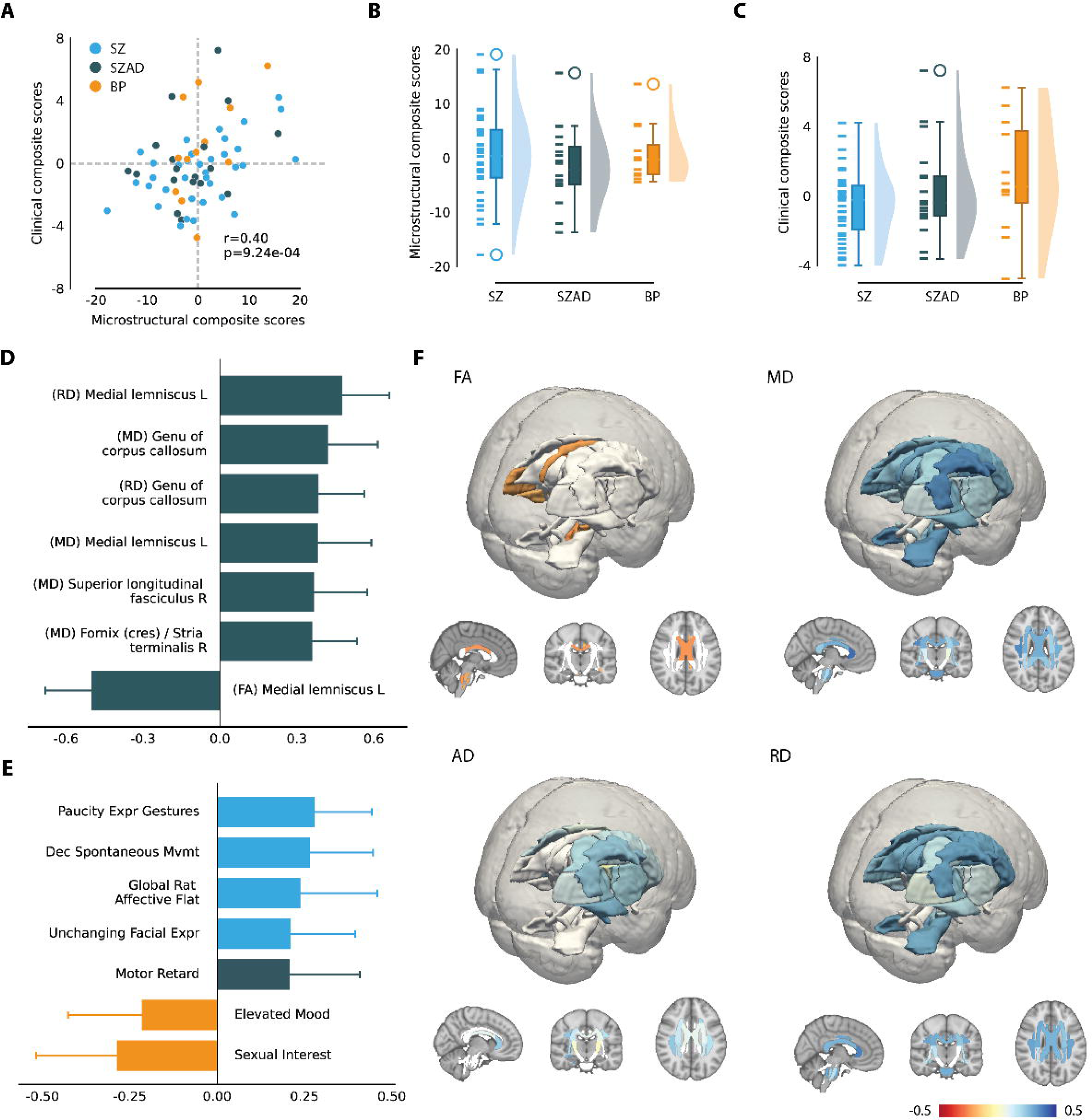
Primary Latent Component (LC) in MEND reflects negative symptom dimension. **(A)** Correlation between individual-specific microstructural and clinical composite scores of patients. Scatterplot was color-coded by each primary diagnostic group. **(B)** Group differences in microstructural composite scores after false discovery rate (FDR) correction. **(C)** Group differences in clinical composite scores. **(D)** Thresholded significant microstructural loadings of the LC. Strong loadings with an effect size greater than 2 standard deviations were shown in the bar chart. Error bars indicate bootstrapped standard deviations. **(E)** Unthresholded significant clinical loadings of the LC. Blue, green, and orange colors indicate the symptom dimensions of negative, general, and mania symptoms, respectively. **(F)** The loading coefficient of all significant microstructural features associated with the LC are projected to their corresponding WM tract location in the brain, categorized by DTI metrics. Non-significant tracts are in white.

### Cross-cohort examination of the symptom profiles

The two independent cohorts, despite employing different symptom scales, demonstrated remarkably consistent clinical profiles that were associated with widespread microstructural alterations. These profiles were primarily characterized by a subset of negative symptoms, particularly diminished emotional expression and reactivity, but notably, also extended to individual general symptoms characterized by somatic manifestations and inversely related to mania symptoms featuring heightened arousal and increased energy levels (Figure 1E; Figure 2E; Table S4). Neither cohort exhibited a statistically significant relationship of microstructure with positive symptoms (p>.05). Figure 3 provides an exhaustive overview of the symptomatology in each cohort’s PLS model.

**Figure 3.**
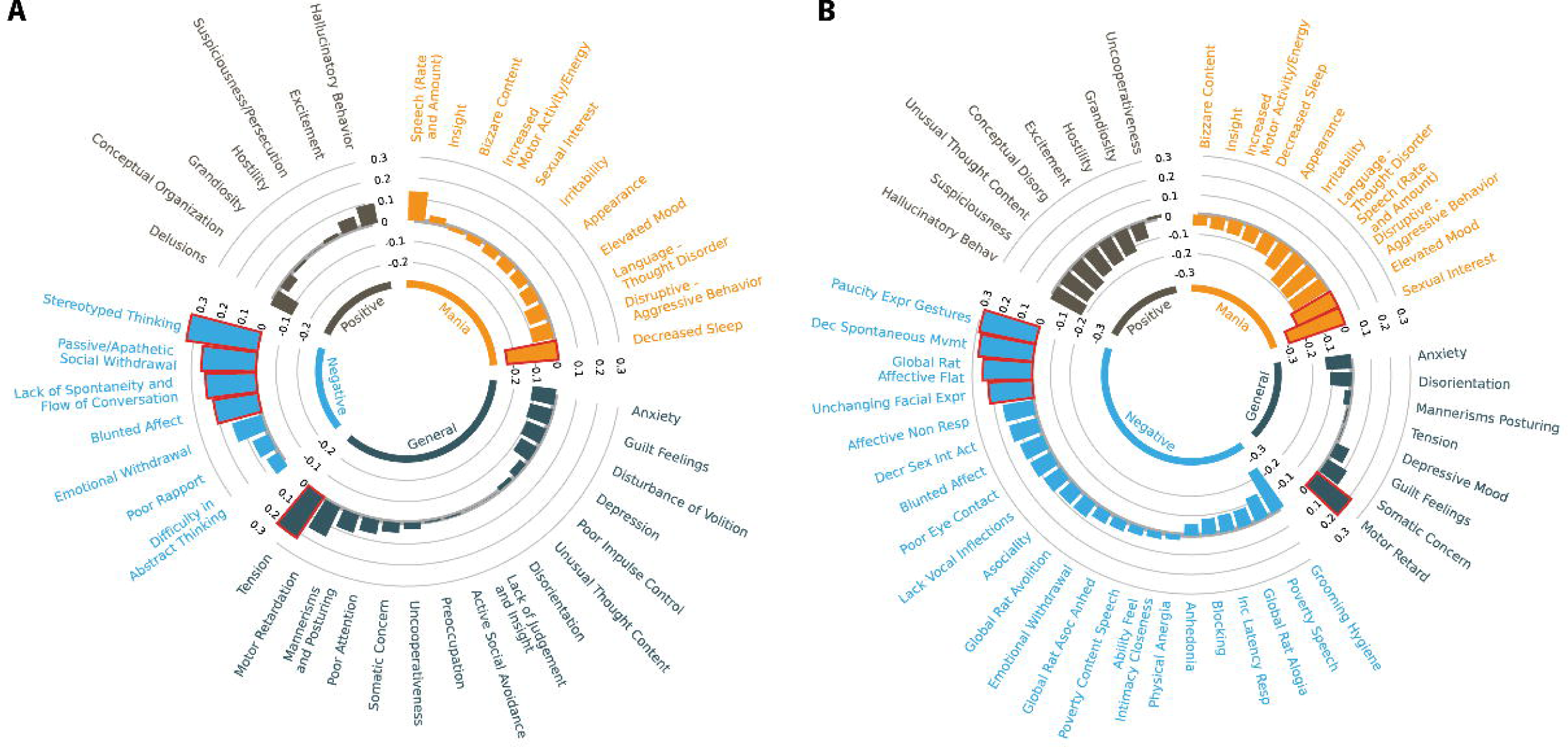
Profiles of multi-dimensional psychopathology corresponding to the identified microstructural feature in each clinical cohort. **(A)** Microstructural features informed dimensions of psychopathology cross clinical rating scales in HCP-EP. **(B**) Microstructural features informed dimensions of psychopathology cross clinical rating scales in MEND. All symptoms reflect the 4 core dimensions of psychopathology. Numbers in the inner rings represent the loading coefficients of symptoms in their respective LC. All loadings in the PLS models are shown here, while red outline of bars indicates loadings with a significant contribution to the higher clinical composite score linking WM alterations in the whole brain (see Figure 1F & 2F).

### Cross-cohort examination of the microstructural signature

To assess the robustness of the identified microstructural signature, we executed a cross-cohort correlation analysis targeting the loading coefficients attributed to each microstructural feature. Our analysis revealed a statistically significant correlation (r=.79, p<.001) between the loadings of the HCP-EP and MEND cohorts, suggesting replicability and generalizability of the observed microstructural signature across independent samples (Figure 4A). Specifically, we noted that features indicative of WM disruptions—namely higher MD, AD, and RD—exhibited a positive correlation with the clinical composite scores, while FA displayed an opposite relationship. For deeper biological insights, we grouped WM tracts into four categories based on their interconnected regions (30) (i.e., brainstem, commissural, association, and projection fibers; Supplemental Methods S10). A Pearson’s correlation conducted on the loading coefficients across both cohorts unveiled a complex landscape of microstructural alterations, distributed across WM tracts of varying connection types (Figure 4B; Figure S3A-E). Additionally, post-hoc tests confirmed that the microstructural signatures were independent of potential confounds, including age, sex, IQ, site, head motion, diagnosis, duration of illness, substance use, or antipsychotic medication, after FDR correction (Table 2).

**Figure 4.**
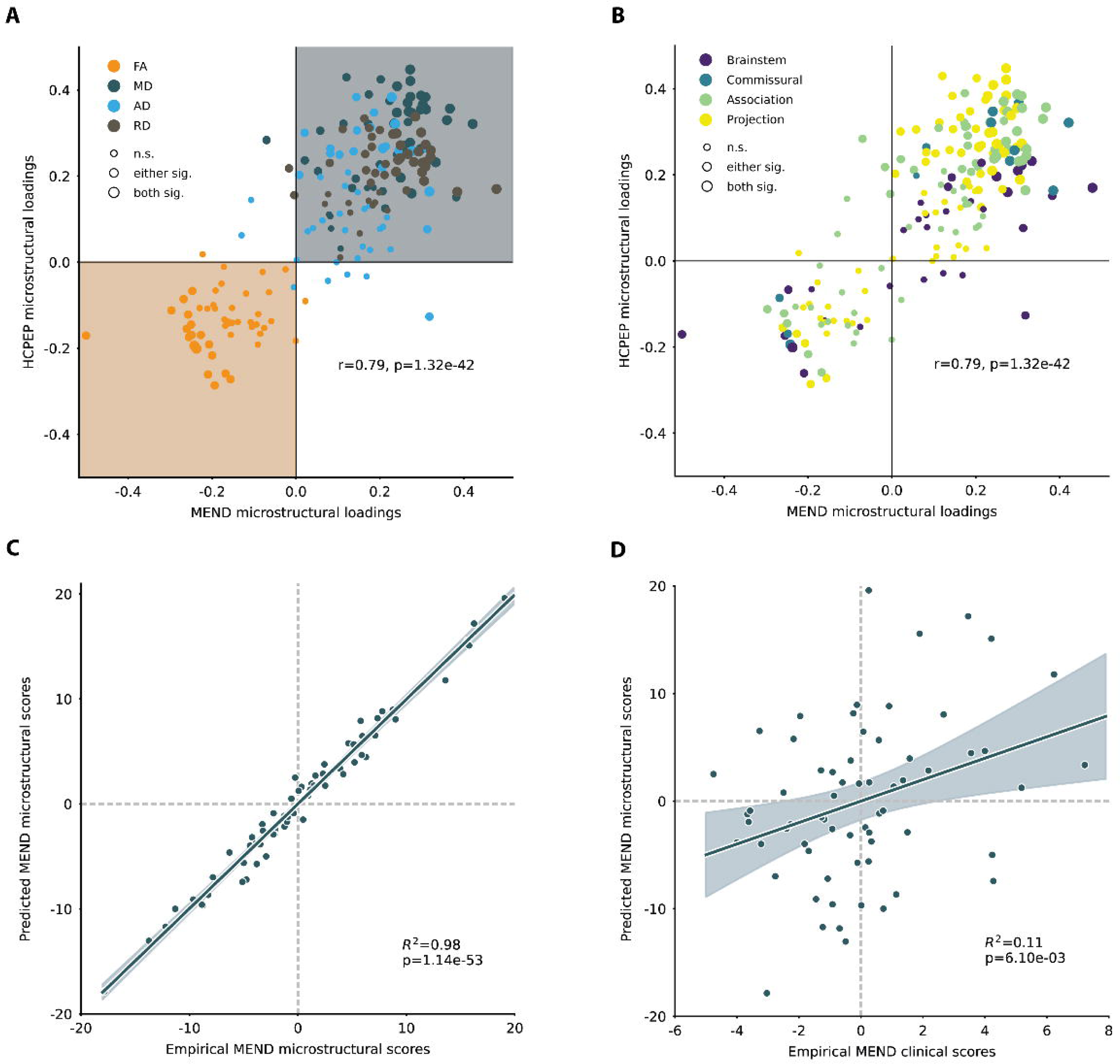
Prediction accuracy of PLS model across independent cohorts. **(A)** Correlation of microstructural features between cohorts, color coded by DTI metrics. **(B)** Correlation of microstructural features between cohorts, color coded by WM tract connectivity types. These groups were defined according to the cortical and subcortical regions interconnected by these tracts, encompassing tracts within the brainstem as well as projection, association, and commissural tracts within the cerebral hemispheres, in alignment with the classification by Oishi (2011) (30). **(C)** Correlation between predicted microstructural composite score and empirical microstructural composite score in the test cohort. **(D)** Correlation between predicted microstructural composite score and empirical clinical composite score in the test cohort.

**Table 2.** Associations between microstructural (or clinical) composite scores and confounds.

|  | HCP -EP |  |  |  | MEND |  |  |  |
| --- | --- | --- | --- | --- | --- | --- | --- | --- |
|  | Microstructural composite score |  | Clinical composite score |  | Microstructural composite score |  | Clinical composite score |  |
|  | *r/t* | *q* | *r/t* | *q* | *r/t* | *q* | *r/t* | *q* |
| Age | 0.00 | 1.00 | 0.00 | 1.00 | 0.00 | 1.00 | 0.00 | 1.00 |
| Sex | 0.00 | 1.00 | 0.00 | 1.00 | 0.00 | 1.00 | 0.00 | 1.00 |
| IQ | 0.00 | 1.00 | 0.00 | 1.00 | 0.00 | 1.00 | 0.00 | 1.00 |
| Site | 0.00 | 1.00 | 0.00 | 1.00 | - | - | - | - |
| Motion | 0.00 | 1.00 | 0.00 | 1.00 | 0.00 | 1.00 | 0.00 | 1.00 |
| Substance use | -0.97 | 0.34 | 0.85 | 0.4 | -0.31 | 0.76 | -0.05 | 0.96 |
| Duration of illness | -0.02 | 0.81 | -0.20 | 0.04* | -0.12 | 0.59 | -0.22 | 0.17 |
| Medication | 0.11 | 0.24 | 0.33 | <.01* | 0.12 | 0.52 | 0.12 | 0.46 |
| Diagnosis<br>(SZ vs BP) | 1.06 | 0.29 | -2.33 | 0.04* | -0.43 | 0.81 | -1.68 | 0.17 |
| Diagnosis<br>(NAP vs AP) | 0.69 | 0.41 | -3.23 | 0.01* | 0.57 | 0.69 | -1.79 | 0.17 |

Either Pearson's correlations (for continuous measures) or t tests (for categorical measures) were computed across all participants. Associations that survived FDR correction ( $q < .05$ ) are indicated with \*.
*Abbreviations: SZ=Schizophrenia spectrum disorders, including schizophrenia, schizoaffective disorder, schizophreniform, psychosis NOS, delusional disorder, or brief psychotic disorder.*
*BP=Bipolar disorders with psychotic features, here for simplicity we also included major depressive disorder with psychotic features. NAP=Non-affective psychotic disorders, including schizophrenia, schizophreniform, psychosis NOS, delusional disorder, or brief psychotic disorder. AP=Affective psychotic disorders, including schizoaffective disorder, bipolar disorders with psychotic features, and major depressive disorder with psychotic features.*

### Cross-cohort prediction

To rigorously assess the generalizability of the identified microstructural signature and its clinical utility, we employed a predictive analysis approach using PLS modeling. The weights assigned to each microstructural feature in the PLS model trained on the HCP-EP cohort were used to compute anticipated microstructural composite scores for each participant in the MEND validation cohort. The projected microstructural composite scores in MEND exhibited a strong correlation with the empirically observed microstructural composite scores (R^2^=.98, p<.001; Figure 4C), evidencing the generalizability and robustness of the microstructural signature across disparate datasets. Further validation was sought by correlating these predicted microstructural composite scores with the empirical clinical composite scores within the MEND cohort. Despite being derived from different clinical scales, a significant albeit moderate correlation was observed (R^2^=.11, p<.001; Figure 4D). This demonstrated the clinical utility of the microstructural signature, holding a significant and specific relationship with psychopathology across independent clinical samples.

### Post-hoc analyses

Finally, we tested whether the microstructural (or clinical) composite scores were associated with confounds, including age, sex, IQ, site, head motion, substance use, duration of illness, medication, and diagnoses. In both cohorts, the microstructural composite scores were not correlated with any confounds after FDR correction (q>.05). In HCP-EP, the clinical composite score was positively correlated with duration of illness (q=.04) and antipsychotic medication dosage (q<.01). Participants with non-affective psychotic disorders, such as schizophrenia, also showed higher clinical composite scores than participants with affective psychotic disorders, such as bipolar disorder with psychotic features (q=.01). In MEND, the clinical composite score was not correlated with any confounds (q>.05), although trending in the same direction as HCP-EP (Table 2).

## Discussion

Using a data-driven, multivariate approach, we found a robust covarying pattern between WM alterations and specific symptom profiles across EP diagnoses. This microstructural signature is correlated with a specific subset of negative symptoms, as well as symptoms related to arousal that extend beyond the conventional scope of the negative symptom category. Notably, this correspondence was confirmed in an independent cohort and the microstructural signature exhibited high predictive accuracy, suggesting replicability and potential clinical utility. Also importantly, exploratory analyses showed the signature was not impacted by antipsychotic medication, substance use, or diagnosis. Together, these findings indicate a central link between brain structural alterations and clinical manifestations within a transdiagnostic framework.

Our findings reinforce previous research in psychotic illnesses by confirming a relationship between WM alterations and clinical symptoms (9). While existing research has primarily been optimized to uncover symptom-to-tract relationships within specific diagnoses (31–34), our analysis, using a statistical-learning method, mapped diagnosis-agnostic relationships between microstructure and psychopathology. One strength of this data-driven approach is its capacity to unveil novel symptom profiles or clusters based on shared neural foundations. Both HCP-EP and MEND cohorts revealed a number of negative symptoms correlating with the neural signature within the dimension of diminished emotion expression (EXP) (35,36). Conversely, only one symptom, apathetic social withdrawal, was within the dimension of motivation and pleasure (MAP) (37–40). Recent confirmatory factor analyses on negative symptoms structure have established five-first-order domains (blunted affect, alogia, anhedonia, avolition, and asociality) in addition to the known second-order dimensions of EXP and MAP (41–44). However, most studies on neurobiological associations of negative symptoms used summarized scores or the broader MAP and EXP dimensions (45,46), missing item-specific neural correlates. The power of this approach is demonstrated by the observed clinical profiles which not only feature symptoms in the EXP dimension, but further extend to incorporate presentations not traditionally seen as negative symptoms, including sleep, tension, and motor slowing.

Negative symptoms are prevalent in EP (47) and persist in up to 70% of psychosis patients post-antipsychotic treatment (48). Moreover, the identified cluster of expressive negative symptoms and hypoarousal, reminiscent of the deficit syndrome seen in psychotic disorders (49), warrants further exploration. This syndrome, characterized by enduring primary negative symptoms, has been associated with WM disruptions (50), poorer responses to treatment, and more severe functional impairment (51). Notably, our approach, which included a broader range of scales compared to the traditional PANSS-based deficit syndrome definition, revealed that additional features like somatic symptoms may share neurobiological underpinnings with typical deficit symptoms. By employing comprehensive clinical measures and data-driven methods, our study offers an expanded conceptualization of this syndrome beyond the conventional criteria (50).

Contrary to previous studies, our results did not detect associations between WM alterations and positive symptoms (47,52–54). One possibility is that negative symptoms may have a stronger and more stable link to WM (55). This suggests different neurobiological bases between positive and negative symptoms, with positive symptoms possibly more tied to other neural features like gray matter alterations (56–58) or functional networks (59–61). In addition, negative symptoms start earlier in the course of psychotic illness, potentially leading to greater variance in younger patients or an association with more pervasive deficits (48,62–64), making their WM signature more readily detectable during EP. Furthermore, other symptom domains might require larger samples to detect links with subtle and widespread microstructural changes, or symptom relationships may become more pronounced as illness progresses.

Our symptom-level analyses shed light on distinct microstructural neural correlates with better granularity than previous individual tract-or symptom-based analyses. As such, our findings did not point to a latent component which emphasized a primary role for any one WM tract. Such outcomes are typical in PLS analyses, which often reveal widespread and subtle brain features that univariate methods are not equipped to detect (65,66). In this context, it is notable that our analysis found MD to capture microstructural changes more widely compared to FA, a finding that corroborates recent research on the elevation of extracellular water in early stages of psychosis. This association suggests that MD may more effectively reflect subtle changes in WM that are indicative of early neurobiological alterations in psychotic disorders.

Microstructural alterations in LC_HCP_ (Figure 1) represented a unique combination of tracts including projection fibers connecting subcortical and cortical regions (corona radiata, internal capsule, external capsule) as well as two long range cortico-cortico association tracts that connect to the frontal lobe (SLF, cingulum). Similarly, in LC_MEND_ there is a combination of subcortical and brainstem connections (medial lemniscus, fornix/stria terminalis) and frontal based tracts (SLF, and genu of corpus callosum). The involvement of additional subcortical tracts not commonly reported in earlier studies may signify their mechanistic roles in EP, such as facilitating neural communication pathways between various brain regions and impacting sensory-motor integration and cognitive functions. These tracts, reflecting recent findings on subcortical volume changes in the HCP-EP dataset (67), are crucial for brain connectivity and efficiency. Taken together, this again highlights the strength of PLS, in which tracts that might typically be of interest in schizophrenia such as the SLF and cingulum (13,68–74), are identified but in addition, a profile of subcortical microstructural changes also emerged.

Variability in this unique combination of tracts could be explored as a potential biomarker for negative symptoms or considered when thinking about new treatment targets. Given the complexity of observed symptom profiles, including symptoms that impact both affective and somatomotor functioning, it is reasonable to assume that a comprehensive microstructural signature would encompass broader WM alterations. Notably, this signature demonstrated high stability and generalizability across independent samples, reinforcing its clinical validity and utility. Interestingly, we observed stronger associations between cross-predicted microstructural measures compared to clinical scores (Figure 4), suggesting that neuroimaging techniques might introduce less variability or “noise” than the measurement of psychiatric symptoms, which can be prone to subjective variability. This discrepancy, highlighted by the cross-cohort prediction, underscores the potential of neuroimaging-based measures as reliable and precise tools in psychiatric research.

### Limitations and conclusions

Given the practical challenges of finding independent EP datasets, the primary and replication cohorts exhibited age differences and used different DWI sequences, which might influence DTI measures and tensor algorithm fitting. However, we replicated the results despite these potential confounds. While we accounted for the impact of anti-psychotic medications, other medications remain unexplored. Additionally, the small sample sizes in our post-hoc diagnostic subgroup analysis raise the risk of type 2 error. Methodologically, we opted for DTI metrics and TBSS over other diffusion approaches to measure WM alterations, guided by their broad applicability and comparability across DWI studies. The widespread use of TBSS inferential metrics throughout the literature offers an advantage for comparison with previous findings and replication, as many existing diffusion MRI datasets include single-shell acquisitions. However, this approach may not capture participant-specific tract variations as effectively as direct DWI tractography, potentially limiting the specificity of our findings (75). Moreover, despite the utility of DTI metrics, their limitations in detailing precise microstructural changes underscore the necessity for advanced methodologies (11). Moving forward, embracing more individualized tractography techniques will be crucial for overcoming these limitations and achieving a richer understanding of WM dynamics.

In conclusion, we identified a strong association between widespread WM alterations and a specific subset of negative symptoms, illuminating a shared central link between brain structural changes and clinical manifestations within a transdiagnostic framework. The microstructural signature, unaffected by diagnosis and medication, further demonstrated robustness and clinical utility through external validation and prediction analyses in an independent cohort. By parsing the neurobiological heterogeneity, treatment strategies could be more precisely tailored, potentially enhancing the effectiveness of early pharmacological (76,77) and psychological (41,78,79) interventions for these neural-related domains of negative symptoms. Future research should prioritize longitudinal studies to trace the development of the microstructural signature identified in EP. These studies could reveal critical insights into the pathophysiological progression and timing of WM alterations, enhancing our understanding of their role in the onset and progression of psychosis. This approach would enable researchers to determine whether these microstructural changes are a precursor to, or a consequence of, the development of psychotic symptoms. Furthermore, by incorporating multimodal sources of variance, such as genetic, environmental, and other neurobiological factors, future studies can work towards a more comprehensive understanding of the complex etiology and pathophysiology of EP.

## Author contributions

Author Contributions: Haley Wang had full access to all of the data in the study and takes responsibility for the integrity of the data and the accuracy of the data analysis.

Study concept and design: HRW, ZQL, KHK. Acquisition, analysis, or interpretation of data: HRW, ZQL, HN, CEH, MBT, PD, CS, TPB, RAM, KHK. Drafting of the manuscript: HRW, KHK. Critical revision of the manuscript for important intellectual content: HRW, ZQL, HN, PKP, TPB, DC, LL, CEB, BM, KHK. Statistical analysis: HRW, ZQL, HN, TPB, KHK. Obtained funding: KHK. Administrative, technical, or material support: ZQL, HN, MBT, PDR, CS, CEB, BM, KHK. Study supervision: HRW, KHK.

## Additional Contributions

Sidhant Chopra, Ph.D., Ariel Rokem, Ph.D., Tiffany C. Ho, Ph.D., Lucina Q. Uddin, Ph.D., Michael F. Green, Ph.D., and Gregory A. Miller, Ph.D. provided critical feedback for the study. Subjects were provided monetary compensation for their participation.

## Acknowledgement

Research using Human Connectome Project for Early Psychosis (HCP-EP) data reported in this publication was supported by the National Institute of Mental Health of the National Institutes of Health under Award Number U01MH109977. The HCP-EP 1.1 Release data used in this report came from DOI: 10.15154/1522899. This work used computational and storage services associated with the Hoffman2 Shared Cluster provided by UCLA Institute for Digital Research and Education’s Research Technology Group. The MEND sample and analysis was supported with NIMH grant MH101506 (KHK) with further support from MH116433 (KHK).

## Data Sharing Statement

All data and code used to perform the analyses can be found at https://github.com/haleyrwang/CCNL_DecodingEP. The HCP-EP dataset is available at https://db.humanconnectome.org/. The MEND dataset is available through the ENIGMA consortium at https://github.com/MICA-MNI/ENIGMA).

