## Supplemental material 1 for "Decoding Early Psychoses: Unraveling Stable Microstructural Features Associated with Psychopathology Across Independent Cohorts"

***Supplemental Information***

**Supplemental Methods**

**Methods S1.** Participant details

**Methods S2.** Differences in symptom severity across diagnostic groups

**Methods S3.** Categorization and dosage of medication

**Methods S4.** Substance use

**Methods S5.** MRI data acquisition

**Methods S6.** Diffusion imaging data analysis

**Methods S7.** Partial least squares analysis details

**Methods S8.** Sensitivity analysis and robustness control

**Methods S9.** Post-hoc analyses on the associations between composite scores and confounds

**Methods S10.** Exploratory analyses of loading coefficients by connectivity type of WM tracts

**Supplemental Results**

**Table S1.** All clinical measures used in the PLS analyses

**Table S2.** Group scores on all clinical measures included in the PLS analyses

**Figure S1.** Differences in symptom severity across diagnostic groups

**Table S3.** Associations between microstructural features and microstructural composite scores

**Table S4.** Associations between clinical features and clinical composite scores

**Figure S2.** Covariance explained by each latent component obtained with the PLS analyses

**Figure S3.** Loading coefficients by connectivity type of WM tracts

**Figure S4.** Tract-specific effects revealed by PLS

**Supplemental References**

This supplementary material has been provided by the authors to give readers additional information about their work.

***Methods S1.*** *Participant details*

Primary cohort: Human Connectome Project for Early Psychosis (HCP-EP, release 1.1)

HCP-EP subjects included 124 EP patients between the ages of 16 and 35 years at study entry. Inclusion criteria for all patients as defined by the HCP included a primary psychotic illness diagnosis (diagnosis of schizophrenia, schizophreniform, schizoaffective, psychosis NOS, delusional disorder, or brief psychotic disorder), or a diagnosis of major depression with psychosis or bipolar disorder with psychosis, based on the Diagnostic and Statistical Manual of Mental Disorders (DSM-5). All patients required a time of psychosis onset within the past 3 years prior to study entry. Exclusion criteria for all participants included IQ less than 70, history of neurological disorders or significant head trauma, and MRI contraindications (e.g., mental implants and braces). Three individuals were excluded from neuroimaging analyses due to missing T1w structural scan. Three individuals were excluded from clinical analyses as they did not complete clinical ratings on psychiatric symptoms. All clinical and cognitive assessments were administered by the HCP team as described in HCP-EP_Release_1.1_Manual. Clinical symptom ratings were evaluated in the EP group using the Positive and Negative Syndrome Scale (PANSS)(1) and Young Mania Rating Scale (YMRS)(2).

Replication cohort: The Multimodal Evaluation of Neural Disorders (MEND)

MEND subjects included 78 EP participants between the ages of 16 and 25 at entry of a longitudinal study at Zucker Hillside Hospital and the Feinstein Institute for Medical Research in New York (MH101506). The study protocol was approved by the Institutional Review Board of Northwell Health. Patients aged 18 and older were provided with written informed consent. Minor participants were provided with written assent along with parental written informed consent. Inclusion criteria for the EP group included duration of illness less than 3 years, current treatment with atypical antipsychotic medication, and a primary psychotic illness diagnosis including Schizophrenia, Schizoaffective, Bipolar Disorder with Psychotic Features, Major Depressive Disorder with Psychotic Features, and Unspecified Psychotic Disorder, based on DSM-5. Exclusion criteria for both all participants included IQ less than 70, insufficient fluency in the English language, and history of neurological disorders or significant head trauma. Clinical symptom ratings were evaluated in the EP group using the Brief Psychiatric Rating Scale (BPRS)(3), the Scale for the Assessment of Negative Symptoms (SANS)(4), and Young Mania Rating Scale (YMRS)(2). A subset of DTI data from the MEND cohort has been previously analysed published in a multi-site mega-analysis study(5).

***Methods S2.*** *Differences in symptom severity across diagnostic groups*

We examined group differences in clinical measures across diagnoses included in the PLS analysis. For the HCP-EP cohort, we compared total scores for PANSS positive symptoms, PANSS negative symptoms, PANSS general symptoms, and YMRS. For the MEND cohort, we compared total scores for BPRS psychotic symptoms, BPRS non-psychotic symptoms, SANS, and YMRS. ANOVA was performed in Python to determine the significance of these clinical differences across diagnostic groups, while controlling for covariates including sex and age (Table S2; Figure S1).

***Methods S3.*** *Categorization and dosage of medication*

In both cohorts, the current dosage of antipsychotic medications was further converted to the equivalent dosage of chlorpromazine. Antipsychotic medication load was tested for association with the microstructural (or clinical) composite scores using Person’s correlations (Table 2). Other types of medication, such as mood stabilizers, antidepressants, sedatives / hypnotics / anxiolytics (SHA), stimulants, were not accounted for in this study due to lack of information from the public datasets.

***Methods S4.*** *Substance use*

Subjects in both cohorts were screened for substance use on the day of scan visit. In HCP-EP cohort, patients with substance use disorders were either mild or in remission. In MEND cohort, patients with current substance use disorders were excluded. In this study, substance use refers to cannabis use in the past 30 days. This choice was motivated by the high prevalence use of cannabis in individuals with psychotic disorders. For both cohorts, cannabis use in the past month was divided into two groups - absent or present - without any further information on the frequency. Substance use was tested for association with the microstructural (or clinical) composite scores using two-sample t test (Table 2).

***Methods S5.*** *MRI data acquisition*

Primary cohort: HCP-EP

Imaging data were collected on three Siemens MAGNETOM Prisma 3T scanners across 4 sites. Scanning sessions included a collection of T1-weighted (multi-echo 3D MPRAGE, TR/TE/TI 2400/2.22/1000ms, 8° flip angle, FOV 256mm, 0.8mm isotropic resolution, 2x GRAPPA acceleration, 208 sagittal slices) and T2-weighted (3D variable-flip-angle turbo-spin-echo (TSE) sequence SPACE, TR/TE 3200/563ms, FOV 256mm, 0.8mm isotropic, 2x GRAPPA, turbo factor 314ms, 208 sagittal slices) structural scans. Additionally, spin echo field maps were collected 4 times with twice for each AP and PA phase encoding (TR/TE 8000/66ms, FoV 208mm, 90° flip angle, 2mm isotropic resolution). Multishell diffusion weighted imaging (DWI) data were collected 4 with both AP and PA phase encoding (TR/TE 3230/89.2ms, FOV 210mm, 1.5mm isotropic resolution, 92 axial slices, b = 1500 and 3000s/mm2 shells, 99/100/108 directions, 15 b0, paired phase encoding).

Replication cohort: MEND

All scans were collected on a 3T Siemens Verio magnetic resonance machine at Zucker Hillside Hospital in Long Island, NY. Scanning sessions include collection of high-resolution T1-weighted (TR/TE 2530/3.3ms, 1mm slice thickness, 7° flip angle, matrix 256*256mm, 1 average) and T2-weighted structural image data (TR/TE 1500/27ms, FOV 220*220mm, 0.5*0.5*3mm isotropic resolution, 1 average) and single shell DWI data (TR/TE 6000/87ms, FOV 240mm, 2.5mm isotropic resolution, 52 axial slices, 65 directions, b = 1000 s/mm2, paired phase encoding).

***Methods S6.*** *Diffusion Imaging Data Analysis*

Raw structural and diffusion MRI data in NIfTI format were acquired from HCP-EP and MEND. Preprocessing and quality control for HCP-EP were performed using HCP Pipelines(6) implemented through Quantitative Neuroimaging Environment & Toolbox (QuNex)(7), while FMRIB Software Library (FSL) v6.1(8) was used for MEND. Specifically, diffusion preprocessing steps for both datasets include intensity normalization, susceptibility distortion correction, eddy current correction, motion correction, and gradient distortion correction(9). Visual and computational quality control procedures were carried out before and after eddy current correction. To estimate DTI metrics indexing WM integrity, dtifit from FSL v6.1 was used to calculate FA, MD, RD, and AD using the b=1500 shell. The maps of the 4 DTI metrics were skeletonized using the Tract-Based Spatial Statistics (TBSS)(10). To increase the generalizability of the results, mean values of DTI metrics were calculated for each diffusion metric along the TBSS skeleton within each WM tract defined by the JHU White-Matter Tractography atlas (JHU-ICBM-DTI-81, distributed with FSL). Additionally, neuroCombat(11) was employed to harmonize multi-site diffusion imaging data to adjust for scanner difference in the HCP-EP sample. In both HCP-EP and MEND, excessive motion in the structural T1w image and diffusion scan image were examined by visual QA on all subjects completing DWI; images were manually categorized as Poor, Good, or Excellent. Following the correction of distortions induced by eddy currents and subject movement, we calculated translation and rotation parameters for each volume through affine registration. Additionally, we assessed slice-by-slice signal drop-out, a DWI-specific issue, using methods outlined by Yendiki et al., 2014 and Benner et al., 2011(12, 13). Subjects with motion parameters exceeding 2 standard deviations were excluded. In the HCP-EP dataset, 3 patients were excluded due to missing T1w images. In MEND, 12 patients were excluded due to motion artifacts.

***Methods S7.*** *Partial least squares analysis*

PLS correlation is an unsupervised algorithm which identifies linear relationships between two sets of features that describe the maximum covariance between them(14). Using the Pyls 0.1.7 Python package(15), we executed PLS correlation between brain microstructural features (4 DTI metrics across 48 WM tracts in the whole brain, n=192) and psychopathology features (n=41 for the HCP-EP cohort and n=49 for the MEND cohort). Briefly, PLS works by decomposing a cross-product matrix (correlation between the microstructural and clinical matrix, a 192x41 (or 192 x 49) matrix in the current study) via singular value decomposition (SVD). The SVD yields 3 output matrices: 2 for microstructural and clinical loadings (i.e., the degree to which that feature contributes to the latent relationship), and a singular value matrix depicting the strength of the latent relationships. The relationship between respective loadings (from the microstructural and clinical matrix) comprises a latent component (LC), representing the latent neuro-psychopathology relationship.

The total number of LCs produced is dictated by the dimensionality of the smaller feature matrix. To better understand the latent neuro-psychopathology relationships at the participant level, the microstructural and clinical loadings from each LC are separately projected onto the respective original microstructural and clinical matrix to create 41 or 49 latent scores (analogous to factor scores). See details of the analytical procedure in a prior study by Nakua et al.(16). Prior to performing the PLS, age, quadratic age, and sex were controlled for by linear regressions, and the residuals were z-transformed.

Feature Contribution Assessment: Each feature's contribution within an LC was gauged using squared-loadings, akin to effect sizes. A z-transformation was applied to these contributions, with a z-score above 2 signifying a feature's significant impact on the LC.

Assessing Significance: To determine the statistical significance of the LCs, we performed a permutation test on the singular values. Specifically, the stability of both the singular values and each feature within a LC (e.g., hallucination) were determined by bootstrap resampling (5,000 repetitions with replacement). A feature was deemed stable in an LC if its 95% confidence interval from the bootstrap distribution excluded zero. To scrutinize the reproducibility of the loadings, a split-half resampling analysis was conducted. In this process, original microstructural and clinical matrices were partitioned into two random halves 10,000 times, and PLS was performed on each. Pearson correlations were then computed between the loadings of each half to measure similarity. Higher correlation coefficients indicated greater reproducibility of loadings across iterations.

Parameter Dimension and Sensitivity: To address the question of whether the validity of the analyses could be affected by the input parameter space or if the model integrity could be improved by reducing the input parameters, we performed additional analyses focusing on the contribution of summary measures (FA and MD) and components (AD and RD) to the microstructural signatures. Specifically, to test if the pattern of results is similar when only the summary measures or components are used, we re-performed the PLS analysis for both cohorts, in 2 different ways: 1) a PLS model with only summary measures (i.e., FA and MD), and 2) a PLS model with only components (i.e., AD and RD). We then correlated the microstructural scores from these 2 new models with the microstructural scores from the original model. Person's correlation analysis showed significant correlations between the summary model and the original model (r=.995, p<.000), and between the component model and the original model (r=.987, p<.000) in HCP-EP. MEND also showed significant correlations between the summary model and the original model (r=.992, p<.000), and between the component model and the original model (r=.988,p<.000). The high correlations indicated that the actual brain pattern corresponding to the clinical features is relatively stable regardless of whether the summary measures are included or not.

***Methods S8.*** *Sensitivity analysis and* *robustness control*

To ensure that the results found were robust, we conducted a series of analyses. First, to bolster the credibility of the PLS result of each sample, we implemented cross-validation through split-sample analysis. Additionally, all loadings underwent bootstrap resampling to enhance stability and verify the reliability of our outcomes. Second, to detect multivariate outliers in the PLS results in both cohorts, we employed the Mahalanobis distance metric, which is particularly effective in identifying outliers in multivariate data sets. Mahalanobis distances did not detect any observations that exceeded 2 standard deviations (SD) for outlier exclusion. Furthermore, we ensured that the application of the neuroCombat did not introduce any confounding effects or alter our results. Pearson’s correlation analysis between PLS results with and without neuroCombat correction yield significantly correlated microstructural composite scores between PLS models (p<.001). To ensure that no phenotype-related WM effect might be attenuated by harmonization, an ANOVA was conducted to assess the impact of site on mean FA values. The analysis revealed that the site was not significantly associated with differences in mean FA values (F_(3, 117)_=.306, p=.821). Lastly, we conducted exploratory analyses of the effects of global DTI metrics and specific tracts. Pearson's correlation coefficients were calculated between the PLS-derived microstructural composite scores, global FA, global MD, and clinical composite scores for each dataset. Fisher's z-transformation was used to compare the strengths of the correlations between the PLS-derived measures, global FA, global MD, and clinical scores (Figure S3A-H).

***Methods S9.*** *Post-hoc analyses on the associations between composite scores and confounds*

We tested whether microstructural and clinical composite scores for LCs of each cohort were associated with sex, age, quadratic age, diagnoses, duration of illness, acquisition site, head motion, substance use, and anti-psychotic medication load. Pearson’s correlations were performed for continuous measures, and t-tests were performed for binary measures (Table 2). For these analyses, all participants from the PLS analyses were included. To address multiple testing, we used the Bonferroni correction method (q<.05) across four families of tests: two each for microstructural and clinical composite scores in the HCP-EP and MEND datasets. Each family had 8 tests corresponding to the 8 confounds.

***Methods S10.*** *Exploratory analyses of loading coefficients by connectivity categories of WM tracts*

To elucidate the biological significance of observed microstructural signatures, we stratified white matter (WM) tracts into four categories, as delineated by the interconnected cortical and subcortical regions. These categories included brainstem tracts, projection tracts, association tracts, and commissural tracts within the cerebral hemispheres, consistent with Oishi's 2011 classification(17). In both the HCP-EP and MEND cohorts, we quantified the density and proportion of each tract category with respect to their loading coefficients (Figure S4A-D). Kernel density analysis was then applied to these loading coefficients across both cohorts, offering a nuanced portrait of microstructural alterations distributed across WM tracts of varying functional connectivity (Figure S4E).

***Table S1.*** *Clinical measures used in the PLS analyses.* This list includes both measures that were available for all participants (and thus included in the PLS analysis) and measures with missing data (marked with *, missing data was handled using median imputation in PLS analysis). The psychopathology dimensions defined in the study for each item in the original clinical scales that are shown below.

***Table S1a.*** *HCP-EP clinical measures used in the PLS analysis.*

| **Scale** | **Subscale/Domain**  (Defined in scale) | **Symptom** | **Dimension**  (Defined in study) | **Missing subjects** |
| --- | --- | --- | --- | --- |
| PANSS | Positive | Delusions | Positive | NA |
|  |  | Conceptual Organization | Positive | NA |
|  |  | Hallucinatory Behavior | Positive | NA |
|  |  | Excitement | Positive | NA |
|  |  | Grandiosity | Positive | NA |
|  |  | Suspiciousness/Persecution | Positive | NA |
|  |  | Hostility | Positive | NA |
|  | Negative | Blunted Affect | Negative | NA |
|  |  | Emotional Withdrawal | Negative | *(2) |
|  |  | Poor Rapport | Negative | *(1) |
|  |  | Passive/Apathetic Social Withdrawal | Negative | *(1) |
|  |  | Difficulty in Abstract Thinking | Negative | *(2) |
|  |  | Lack of Spontaneity and Flow of Conversation | Negative | *(1) |
|  |  | Stereotyped Thinking | Negative | *(1) |
|  | General | Somatic Concern | General | NA |
|  |  | Anxiety | General | *(1) |
|  |  | Guilt Feelings | General | *(1) |
|  |  | Tension | General | NA |
|  |  | Mannerisms and Posturing | General | *(1) |
|  |  | Depression | General | *(1) |
|  |  | Motor Retardation | General | NA |
|  |  | Uncooperativeness | General | NA |
|  |  | Unusual Thought Content | General | *(1) |
|  |  | Disorientation | General | *(1) |
|  |  | Poor Attention | General | *(1) |
|  |  | Lack of Judgement and Insight | General | *(1) |
|  |  | Disturbance of Volition | General | *(1) |
|  |  | Poor Impulse Control | General | NA |
|  |  | Preoccupation | General | *(1) |
|  |  | Active Social Avoidance | General | *(2) |
| YMRS | Mania | Elevated Mood | Mania | NA |
|  |  | Increased Motor Activity-Energy | Mania | NA |
|  |  | Irritability | Mania | NA |
|  |  | Language-Thought Disorder | Mania | NA |
|  |  | Content | Mania | NA |
|  |  | Disruptive-Aggressive Behavior | Mania | NA |
|  |  | Appearance | Mania | NA |
|  |  | Insight | Mania | NA |
|  |  | Increased Speech (Rate and Amount) | Mania | NA |
|  |  | Increased Sexual Interest | Mania | NA |
|  |  | Sleep | Mania | NA |

***Table S1b.*** *MEND* *clinical measures used in the PLS analysis.*

| **Scale** | **Subscale/Domain**  (Defined in scale) | **Symptom** | **Dimension**  (Defined in study) | **Missing subjects** |
| --- | --- | --- | --- | --- |
| BPRS | Psychotic | Conceptual Disorganization | Positive | NA |
|  |  | Grandiosity | Positive | NA |
|  |  | Hostility | Positive | NA |
|  |  | Suspiciousness | Positive | NA |
|  |  | Hallucinatory Behavior | Positive | NA |
|  |  | Uncooperativeness | Positive | NA |
|  |  | Unusual Thought Content | Positive | NA |
|  |  | Excitement | Positive | NA |
|  | Non-psychotic | Anxiety | General | NA |
|  |  | Disorientation | General | NA |
|  |  | Guilt Feelings | General | NA |
|  |  | Tension | General | NA |
|  |  | Mannerisms And Posturing | General | NA |
|  |  | Depressive Mood | General | NA |
|  |  | Motor Retardation | General | NA |
|  |  | Somatic Concern | General | NA |
|  |  | Blunted Affect | Negative | NA |
|  |  | Emotional Withdrawal | Negative | NA |
| SANS | Affective Flattening or Blunting | Unchanging Facial Expression | Negative | NA |
|  |  | Decreased Spontaneous Movements | Negative | NA |
|  |  | Paucity of Expressive Gestures | Negative | NA |
|  |  | Poor Eye Contact | Negative | NA |
|  |  | Affective Nonresponsivity | Negative | NA |
|  |  | Lack of Vocal Inflections | Negative | NA |
|  |  | Global Rating of Affective Flattening | Negative | NA |
|  | Alogia | Poverty of Speech | Negative | NA |
|  |  | Poverty of Content of Speech | Negative | NA |
|  |  | Blocking | Negative | NA |
|  |  | Increased Latency of Response | Negative | NA |
|  |  | Global Rating of Alogia | Negative | NA |
|  | Avolition/  Apathy | Grooming and Hygiene | Negative | NA |
|  |  | Physical Anergia | Negative | NA |
|  |  | Global Rating of Avolition/Apathy | Negative | NA |
|  |  | Asociality | Negative | NA |
|  |  | Decrease in Recreational Interests and Activities | Negative | NA |
|  |  | Decrease in Sexual Interest and Activity | Negative | NA |
|  |  | Ability to Feel Intimacy and Closeness | Negative | NA |
|  |  | Global Rating of Anhedonia/Asociality | Negative | NA |
| YMRS | Mania | Elevated Mood | Mania | NA |
|  |  | Increased Motor Activity-Energy | Mania | NA |
|  |  | Irritability | Mania | NA |
|  |  | Language-Thought Disorder | Mania | NA |
|  |  | Content | Mania | NA |
|  |  | Disruptive-Aggressive Behavior | Mania | NA |
|  |  | Appearance | Mania | NA |
|  |  | Insight | Mania | NA |
|  |  | Increased Speech (Rate and Amount) | Mania | NA |
|  |  | Increased Sexual Interest | Mania | NA |
|  |  | Sleep | Mania | NA |

***Table S2.*** *Differences in symptom severity across diagnostic groups.* Mean (SD) values of the summarized clinical scores are shown for each primary diagnostic group. Groups were compared with ANCOVAs with age and sex as covariates. All p values that survived FDR correction (q<.05) are indicated with asterisks (*=q<.05).

***Table S2a. HCP-EP***

| Scale / Subscale | | SZ | SZAD | BD | F | p value |
| --- | --- | --- | --- | --- | --- | --- |
| PANSS | Positive | 11.54 (3.77) | 14 (4.9) | 8.89 (2.47) | 10.19 | <.001* |
|  | Negative | 15.3 (5.77) | 12.71 (3.27) | 10.96 (3.72) | 6.05 | <.01* |
|  | General | 25.43 (4.94) | 25.64 (6.25) | 22 (4.87) | 4.35 | 0.02* |
| YMRS | Mania | 4.5 (4.52) | 6.79 (8.42) | 4.98 (5.39) | 1.07 | 0.35 |

***Table S2b. MEND***

| Scale / Subscale | | SZ | SZAD | BD | F | p value |
| --- | --- | --- | --- | --- | --- | --- |
| BPRS | Psychotic | 13.51 (4.58) | 13.52 (5.16) | 15.63 (8.39) | 1.51 | 0.23 |
|  | Non-psychotic | 16.77 (4.74) | 18.43 (6.12) | 17.13 (6.28) | 1.08 | 0.34 |
| SANS | Negative | 21.18 (14.53) | 19.52 (11.76) | 12.81 (12.44) | 1.44 | 0.24 |
| YMRS | Mania | 4.72 (4.76) | 7.33 (8.42) | 10.5 (9.7) | 5.51 | <.01* |

*Abbreviations:* *SZ=schizophrenia spectrum disorders, including schizophrenia, schizophreniform, psychosis NOS, delusional disorder, or brief psychotic disorder. SZAD=schizoaffective disorder; BP= bipolar disorders with psychotic features or major depressive disorder with psychotic features.*

***Figure S1.*** *Differences in symptom severity across diagnostic groups.* In HCP-EP, among the diagnostic groups of schizophrenia, schizoaffective disorder, and psychotic mood disorders, significant differences in severity were observed for positive, negative, and general symptoms post FDR correction. We did not find a group difference of mania symptoms (see Table S2 for ANCOVA significances).

***Figure S1A.*** *HCP-EP*

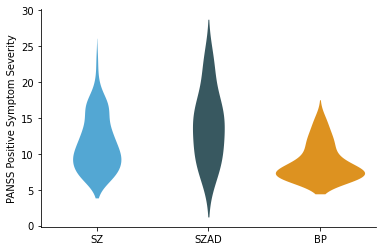

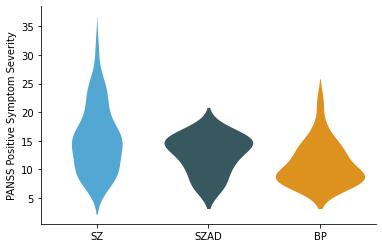
**A	B**

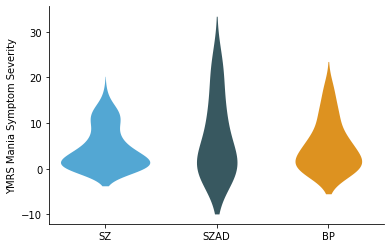
**C	D**

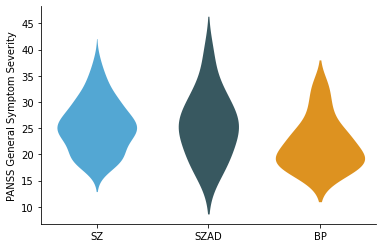

***Figure S1B.*** *MEND*

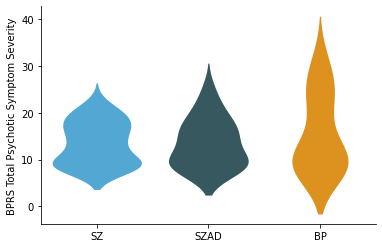

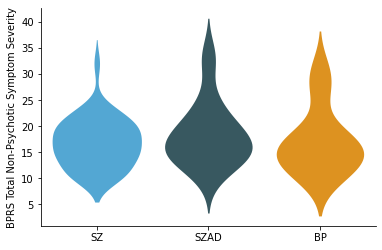
**A	B**

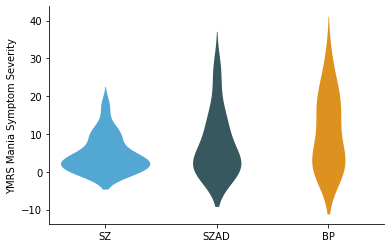

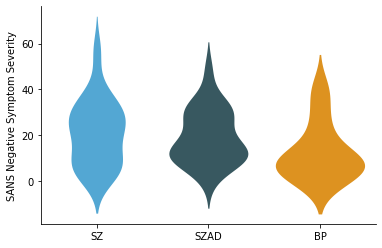
**C	D**

***Table S3.*** *Pearson’s correlations between participants’ microstructural features and their microstructural composite scores.* These correlations show the contribution of each microstructural measure to the identified latent component, along with their bootstrap-estimated standard errors (SE). Features were sorted by decreasing strength (i.e., absolute loading coefficient). Top 50 significant DTI features involved in the latent component are shown in the tables (p<.05). The full list can be found at: <https://github.com/haleyrwang/CCNL_DecodingEP>.

***Table S3A.*** *HCP-EP*

| **DTI metric** | **Tract name** | **Connectivity category** | **Corr (SE)** |
| --- | --- | --- | --- |
| MD | Posterior corona radiata (L) | Projection tracts | 0.45 (0.18) |
| MD | Posterior limb of internal capsule (R) | Projection tracts | 0.43 (0.14) |
| MD | Posterior corona radiata (R) | Projection tracts | 0.42 (0.20) |
| MD | Retrolenticular part of internal capsule (R) | Projection tracts | 0.42 (0.16) |
| MD | Retrolenticular part of internal capsule (L) | Projection tracts | 0.41 (0.14) |
| MD | External capsule (R) | Association tracts | 0.39 (0.17) |
| MD | Superior longitudinal fasciculus (L) | Association tracts | 0.39 (0.18) |
| MD | Anterior corona radiata (R) | Projection tracts | 0.38 (0.16) |
| MD | Cingulum (cingulate gyrus) (L) | Association tracts | 0.38 (0.17) |
| AD | Posterior corona radiata (L) | Projection tracts | 0.38 (0.18) |
| AD | Retrolenticular part of internal capsule (R) | Projection tracts | 0.38 (0.22) |
| MD | Superior corona radiata (L) | Projection tracts | 0.37 (0.17) |
| MD | Sagittal stratum (include inferior longitidinal fasciculus and inferior fronto-occipital fasciculus) (R) | Association tracts | 0.37 (0.19) |
| MD | Body of corpus callosum | Commissural tracts | 0.37 (0.20) |
| MD | Superior longitudinal fasciculus (R) | Association tracts | 0.36 (0.19) |
| MD | Cingulum (cingulate gyrus) (R) | Association tracts | 0.36 (0.19) |
| MD | Superior corona radiata (R) | Projection tracts | 0.36 (0.20) |
| MD | Anterior corona radiata (L) | Projection tracts | 0.35 (0.16) |
| AD | Posterior corona radiata (R) | Projection tracts | 0.35 (0.22) |
| MD | Splenium of corpus callosum | Commissural tracts | 0.35 (0.22) |
| MD | Superior fronto-occipital fasciculus (could be a part of anterior internal capsule) (R) | Association tracts | 0.35 (0.20) |
| RD | Posterior corona radiata (L) | Projection tracts | 0.34 (0.19) |
| RD | Posterior corona radiata (R) | Projection tracts | 0.33 (0.19) |
| MD | Fornix (cres) / Stria terminalis (can not be resolved with current resolution) (R) | Association tracts | 0.33 (0.19) |
| MD | Cingulum (hippocampus) (R) | Association tracts | 0.33 (0.17) |
| MD | Fornix (cres) / Stria terminalis (can not be resolved with current resolution) (L) | Association tracts | 0.32 (0.17) |
| MD | Posterior thalamic radiation (include optic radiation) (R) | Projection tracts | 0.32 (0.16) |
| MD | Sagittal stratum (include inferior longitidinal fasciculus and inferior fronto-occipital fasciculus) (L) | Association tracts | 0.32 (0.16) |
| AD | Genu of corpus callosum | Commissural tracts | 0.32 (0.20) |
| MD | Genu of corpus callosum | Commissural tracts | 0.32 (0.18) |
| AD | Superior longitudinal fasciculus (L) | Association tracts | 0.31 (0.21) |
| RD | Sagittal stratum (include inferior longitidinal fasciculus and inferior fronto-occipital fasciculus) (L) | Association tracts | 0.31 (0.17) |
| RD | External capsule (R) | Association tracts | 0.31 (0.20) |
| RD | Retrolenticular part of internal capsule (L) | Projection tracts | 0.31 (0.14) |
| AD | Superior corona radiata (R) | Projection tracts | 0.31 (0.24) |
| MD | Posterior limb of internal capsule (L) | Projection tracts | 0.31 (0.16) |
| RD | Cingulum (cingulate gyrus) (L) | Association tracts | 0.30 (0.16) |
| AD | Posterior limb of internal capsule (R) | Projection tracts | 0.30 (0.17) |
| RD | Posterior thalamic radiation (include optic radiation) (L) | Projection tracts | 0.30 (0.19) |
| RD | Anterior limb of internal capsule (R) | Projection tracts | 0.30 (0.14) |
| RD | Sagittal stratum (include inferior longitidinal fasciculus and inferior fronto-occipital fasciculus) (R) | Association tracts | 0.29 (0.21) |
| FA | Posterior thalamic radiation (include optic radiation) (L) | Projection tracts | -0.29 (0.21) |
| MD | Tapetum (R) | Association tracts | 0.28 (0.18) |
| MD | Superior fronto-occipital fasciculus (could be a part of anterior internal capsule) (L) | Association tracts | 0.28 (0.16) |
| RD | Anterior corona radiata (R) | Projection tracts | 0.27 (0.18) |
| FA | Anterior limb of internal capsule (R) | Projection tracts | -0.27 (0.15) |
| RD | Superior corona radiata (L) | Projection tracts | 0.27 (0.17) |
| MD | Anterior limb of internal capsule (R) | Projection tracts | 0.27 (0.14) |
| AD | Retrolenticular part of internal capsule (L) | Projection tracts | 0.27 (0.18) |
| RD | Fornix (cres) / Stria terminalis (can not be resolved with current resolution) (L) | Association tracts | 0.27 (0.20) |

***Table S3B.*** *MEND*

| **DTI metric** | **Tract name** | **Connectivity category** | **Corr (SE)** |
| --- | --- | --- | --- |
| FA | Medial lemniscus (L) | Tracts in the brainstem | -0.50 (0.18) |
| RD | Medial lemniscus (L) | Tracts in the brainstem | 0.48 (0.18) |
| MD | Genu of corpus callosum | Commissural tracts | 0.42 (0.19) |
| RD | Genu of corpus callosum | Commissural tracts | 0.38 (0.18) |
| MD | Medial lemniscus (L) | Tracts in the brainstem | 0.38 (0.21) |
| MD | Superior longitudinal fasciculus (R) | Association tracts | 0.37 (0.21) |
| MD | Fornix (cres) / Stria terminalis (can not be resolved with current resolution) (R) | Association tracts | 0.36 (0.18) |
| MD | Middle cerebellar peduncle | Tracts in the brainstem | 0.33 (0.20) |
| RD | Superior longitudinal fasciculus (L) | Association tracts | 0.32 (0.21) |
| RD | Superior longitudinal fasciculus (R) | Association tracts | 0.32 (0.23) |
| AD | Pontine crossing tract (a part of MCP) | Tracts in the brainstem | 0.32 (0.22) |
| AD | Fornix (cres) / Stria terminalis (can not be resolved with current resolution) (R) | Association tracts | 0.32 (0.21) |
| AD | Cingulum (hippocampus) (R) | Association tracts | 0.31 (0.26) |
| AD | Middle cerebellar peduncle | Tracts in the brainstem | 0.31 (0.21) |
| RD | Cingulum (cingulate gyrus) (R) | Association tracts | 0.31 (0.24) |
| MD | Cingulum (cingulate gyrus) (R) | Association tracts | 0.31 (0.26) |
| RD | Anterior corona radiata (R) | Projection tracts | 0.31 (0.23) |
| MD | Cingulum (cingulate gyrus) (L) | Association tracts | 0.31 (0.22) |
| RD | Medial lemniscus (R) | Tracts in the brainstem | 0.31 (0.21) |
| RD | Superior corona radiata (R) | Projection tracts | 0.31 (0.21) |
| MD | Body of corpus callosum | Commissural tracts | 0.30 (0.19) |
| RD | Superior fronto-occipital fasciculus (could be a part of anterior internal capsule) (R) | Association tracts | 0.30 (0.23) |
| RD | Middle cerebellar peduncle | Tracts in the brainstem | 0.30 (0.19) |
| RD | Cingulum (cingulate gyrus) (L) | Association tracts | 0.30 (0.20) |
| MD | Superior longitudinal fasciculus (L) | Association tracts | 0.30 (0.21) |
| MD | Superior corona radiata (R) | Projection tracts | 0.30 (0.22) |
| MD | Posterior thalamic radiation (include optic radiation) (L) | Projection tracts | 0.30 (0.21) |
| FA | Superior longitudinal fasciculus (L) | Association tracts | -0.30 (0.23) |
| RD | Body of corpus callosum | Commissural tracts | 0.29 (0.21) |
| RD | Posterior corona radiata (L) | Projection tracts | 0.28 (0.19) |
| RD | Splenium of corpus callosum | Commissural tracts | 0.28 (0.17) |
| AD | Sagittal stratum (include inferior longitidinal fasciculus and inferior fronto-occipital fasciculus) (R) | Association tracts | 0.28 (0.24) |
| MD | Pontine crossing tract (a part of MCP) | Tracts in the brainstem | 0.28 (0.23) |
| MD | Superior fronto-occipital fasciculus (could be a part of anterior internal capsule) (R) | Association tracts | 0.28 (0.24) |
| MD | Anterior corona radiata (R) | Projection tracts | 0.27 (0.21) |
| MD | Retrolenticular part of internal capsule (R) | Projection tracts | 0.27 (0.18) |
| MD | Posterior corona radiata (L) | Projection tracts | 0.27 (0.20) |
| MD | Superior fronto-occipital fasciculus (could be a part of anterior internal capsule) (L) | Association tracts | 0.27 (0.23) |
| MD | Anterior corona radiata (L) | Projection tracts | 0.27 (0.20) |
| FA | Genu of corpus callosum | Commissural tracts | -0.27 (0.18) |
| RD | Anterior corona radiata (L) | Projection tracts | 0.27 (0.20) |
| MD | Sagittal stratum (include inferior longitidinal fasciculus and inferior fronto-occipital fasciculus) (R) | Association tracts | 0.27 (0.22) |
| RD | Cerebral peduncle (R) | Projection tracts | 0.26 (0.18) |
| FA | Posterior corona radiata (L) | Projection tracts | -0.26 (0.23) |
| FA | Cingulum (cingulate gyrus) (R) | Association tracts | -0.26 (0.21) |
| MD | Anterior limb of internal capsule (R) | Projection tracts | 0.26 (0.15) |
| FA | Middle cerebellar peduncle | Tracts in the brainstem | -0.26 (0.18) |
| FA | Tapetum (L) | Association tracts | -0.25 (0.22) |
| RD | Superior corona radiata (L) | Projection tracts | 0.25 (0.22) |
| MD | Medial lemniscus (R) | Tracts in the brainstem | 0.25 (0.21) |

***Table S4.*** *Pearson’s correlations between participants’ clinical measures and their clinical composite scores.* These correlations show the contribution of each clinical measure to the identified latent component, along with their bootstrap-estimated standard errors (SE). Correlations with significant bootstrapped Z scores (p<.05) are indicated with *.

***Table S4a. HCP-EP***

| **Scale** | **Dimension** (Defined in study) | **Feature** | **Corr (SE)** |
| --- | --- | --- | --- |
| PANSS | Positive | Delusions | -0.11 (0.17) |
|  |  | Conceptual Organization | -0.04 (0.16) |
|  |  | Hallucinatory Behavior | 0.09 (0.19) |
|  |  | Excitement | 0.06 (0.19) |
|  |  | Grandiosity | -0.01 (0.20) |
|  |  | Suspiciousness/Persecution | 0.02 (0.17) |
|  |  | Hostility | 0.00 (0.20) |
|  | Negative | Blunted Affect | 0.20 (0.20)* |
|  |  | Emotional Withdrawal | 0.14 (0.19) |
|  |  | Poor Rapport | 0.08 (0.19) |
|  |  | Passive/Apathetic Social Withdrawal | 0.24 (0.18)* |
|  |  | Difficulty in Abstract Thinking | 0.06 (0.20) |
|  |  | Lack of Spontaneity and Flow of Conversation | 0.23 (0.19)* |
|  |  | Stereotyped Thinking | 0.33 (0.20)* |
|  | General | Somatic Concern | 0.05 (0.22) |
|  |  | Anxiety | -0.11 (0.16) |
|  |  | Guilt Feelings | -0.11 (0.21) |
|  |  | Tension | 0.22 (0.18)* |
|  |  | Mannerisms and Posturing | 0.09 (0.18) |
|  |  | Depression | -0.09 (0.18) |
|  |  | Motor Retardation | 0.15 (0.20) |
|  |  | Uncooperativeness | 0.03 (0.17) |
|  |  | Unusual Thought Content | -0.02 (0.20) |
|  |  | Disorientation | 0.00 (0.18) |
|  |  | Poor Attention | 0.07 (0.17) |
|  |  | Lack of Judgement and Insight | 0.00 (0.19) |
|  |  | Disturbance of Volition | -0.10 (0.18) |
|  |  | Poor Impulse Control | -0.04 (0.17) |
|  |  | Preoccupation | 0.01 (0.20) |
|  |  | Active Social Avoidance | 0.01 (0.16) |
| YMRS | Mania | Elevated Mood | -0.07 (0.11) |
|  |  | Increased Motor Activity-Energy | -0.03 (0.12) |
|  |  | Irritability | -0.05 (0.15) |
|  |  | Language-Thought Disorder | -0.09 (0.14) |
|  |  | Content | -0.02 (0.18) |
|  |  | Disruptive-Aggressive Behavior | -0.10 (0.17) |
|  |  | Appearance | -0.06 (0.15) |
|  |  | Insight | 0.03 (0.18) |
|  |  | Increased Speech (Rate and Amount) | 0.14 (0.24) |
|  |  | Increased Sexual Interest | -0.05 (0.19) |
|  |  | Decreased Sleep | -0.23 (0.10)* |

***Table S4b. MEND***

| **Scale** | **Dimension** (Defined in study) | **Feature** | **Corr (SE)** |
| --- | --- | --- | --- |
| BPRS | Positive | Conceptual Disorganization | 0.15 (0.28) |
|  |  | Grandiosity | 0.09 (0.21) |
|  |  | Hostility | 0.13 (0.23) |
|  |  | Suspiciousness | 0.17 (0.30) |
|  |  | Hallucinatory Behavior | 0.18 (0.28) |
|  |  | Uncooperativeness | -0.02 (0.09) |
|  |  | Unusual Thought Content | 0.17 (0.25) |
|  |  | Excitement | 0.14 (0.21) |
|  | General | Anxiety | 0.13 (0.26) |
|  |  | Disorientation | 0.11 (0.28) |
|  |  | Guilt Feelings | -0.07 (0.29) |
|  |  | Tension | 0.01 (0.21) |
|  |  | Mannerisms And Posturing | 0.03 (0.18) |
|  |  | Depressive Mood | 0.00 (0.29) |
|  |  | Motor Retardation | -0.21 (0.20)* |
|  |  | Somatic Concern | -0.11 (0.27) |
|  | Negative | Blunted Affect | -0.13 (0.23) |
|  |  | Emotional Withdrawal | -0.06 (0.23) |
| SANS | Negative | Unchanging Facial Expression | -0.21 (0.18)* |
|  |  | Decreased Spontaneous Movements | -0.27 (0.18)* |
|  |  | Paucity of Expressive Gestures | -0.28 (0.16)* |
|  |  | Poor Eye Contact | -0.12 (0.20) |
|  |  | Affective Nonresponsivity | -0.16 (0.19) |
|  |  | Lack of Vocal Inflections | -0.10 (0.24) |
|  |  | Global Rating of Affective Flattening | -0.24 (0.22)* |
|  |  | Poverty of Speech | 0.13 (0.21) |
|  |  | Poverty of Content of Speech | -0.04 (0.22) |
|  |  | Blocking | 0.08 (0.16) |
|  |  | Increased Latency of Response | 0.09 (0.22) |
|  |  | Global Rating of Alogia | 0.09 (0.23) |
|  |  | Grooming and Hygiene | 0.24 (0.25) |
|  |  | Physical Anergia | -0.03 (0.25) |
|  |  | Global Rating of Avolition/Apathy | -0.09 (0.24) |
|  |  | Asociality | -0.09 (0.25) |
|  |  | Recreational Interests and Activities | 0.06 (0.25) |
|  |  | Decrease in Sexual Interest and Activity | -0.15 (0.24) |
|  |  | Ability to Feel Intimacy and Closeness | -0.03 (0.27) |
|  |  | Global Rating of Anhedonia/Asociality | -0.05 (0.25) |
| YMRS | Mania | Elevated Mood | 0.22 (0.21)* |
|  |  | Increased Motor Activity-Energy | 0.07 (0.22) |
|  |  | Irritability | 0.14 (0.25) |
|  |  | Language-Thought Disorder | 0.19 (0.27) |
|  |  | Content | 0.05 (0.21) |
|  |  | Disruptive-Aggressive Behavior | 0.20 (0.23) |
|  |  | Appearance | 0.10 (0.26) |
|  |  | Insight | 0.06 (0.22) |
|  |  | Increased Speech (Rate and Amount) | 0.19 (0.23) |
|  |  | Increased Sexual Interest | 0.29 (0.23)* |
|  |  | Decreased Sleep | 0.08 (0.18) |

***Figure S2.*** *Covariance explained by each latent component obtained with the PLS analyses.* The first component (in red) in each cohort were found to be statistically significant by permutation testing with FDR correction (q<.05).

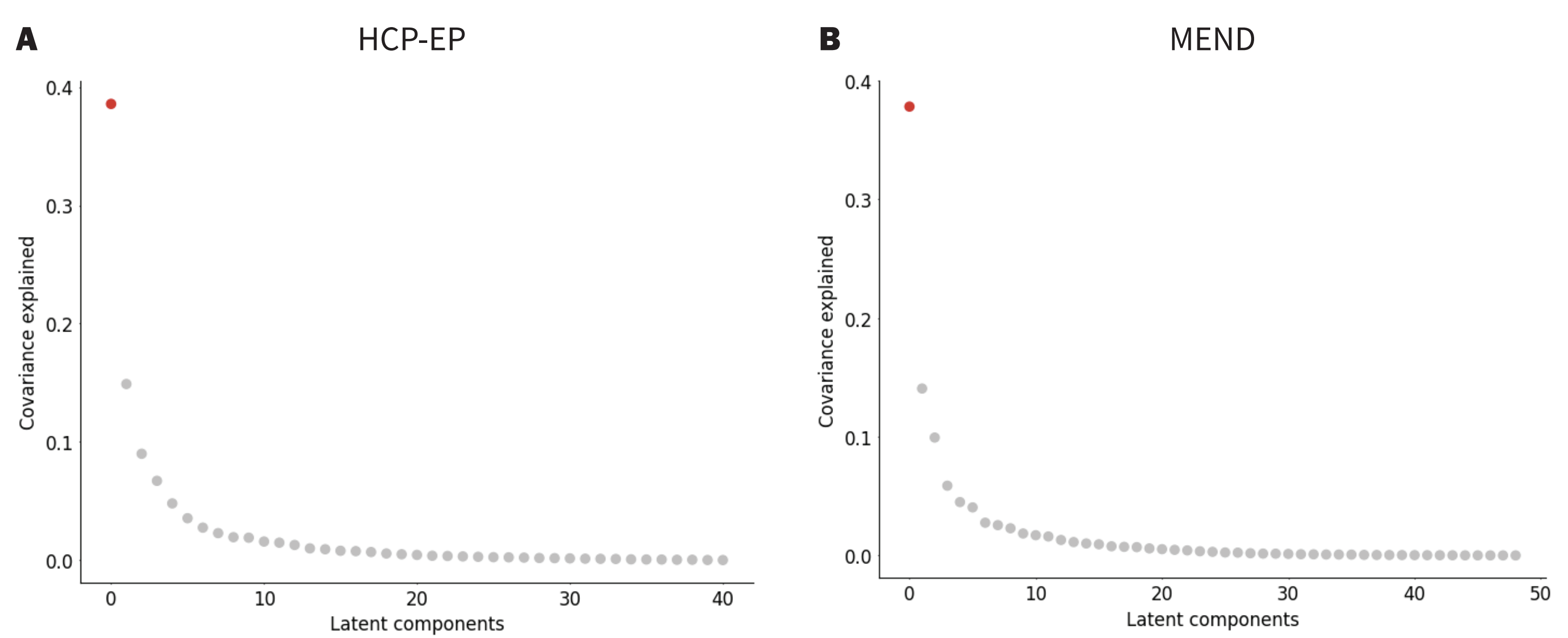

***Figure S3.*** *Tract-specific effects revealed by PLS.* In HCP-EP, the microstructural composite scores were significantly correlated with both global FA (r=-.646, p<.000, Figure S3A) and global MD (r=.869, p<.000, Figure S3B). LC_HCP_ clinical composite scores were also significantly correlated with global FA (r=-.258, p=.004, Figure S3C). and global MD (r=.429, p<.000, Figure S3D). Fisher's z-transformation was applied to assess whether the relationships between LC_HCP_ clinical composite scores with LC_HCP_ microstructural composite scores (r=.435; Figure 1A) and global FA (r=-.258) differed significantly. The transformation showed that the strength of the association between LC_HCP_ clinical scores and LC_HCP_ brain measures is significantly stronger than its association with global FA (z=5.61, p<.000). The correlations between LC_HCP_ clinical scores and LC_HCP_ brain scores are not significantly different from its association with global MD (z=.058, p=.954).

Similarly, in MEND, the microstructural composite scores were significantly correlated with both global FA (r=-.822, p<.000, Figure S3E) and global MD (r=.845, p<.000, Figure S3F). LC_MEND_ clinical composite scores were also significantly correlated with global FA (r=-.299, p=.016, Figure S3G). and global MD (r=.335, p=.006, Figure S3H). Fisher's z-transformation was applied to assess whether the relationships between LC_MEND_ clinical composite scores with LC_MEND_ microstructural composite scores (r=.401; Figure 2A) and global FA (r=-.299) differed significantly. The transformation showed that the strength of the association between LC_MEND_ clinical scores and LC_MEND_ brain measures is significantly stronger than its association with global FA (z=4.08, p<.000). The correlations between LC_MEND_ clinical scores and LC_MEND_ brain scores are not significantly different from its association with global MD (z=.429, p=.668).

Taken together, these findings indicate that global MD is an informative predictor of clinical measures, demonstrating notable correlations with symptom profiles. While the tract-based PLS model did not significantly outperform global MD in terms of its association with clinical scores, it showed significantly stronger correlations with clinical scores than global FA, and the consistently higher correlation coefficients observed between the PLS-derived measures and clinical scores across both datasets suggest that the tract-specific information captured by PLS may offer complementary insights into brain-symptom relationships. Future research should further investigate the unique contributions of tract-specific and global measures in understanding the neurobiological underpinnings of psychopathology, as these approaches may provide distinct yet valuable perspectives on the brain-behavior relationship in EP.

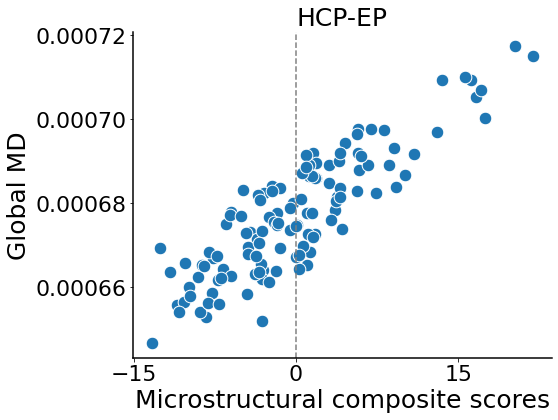

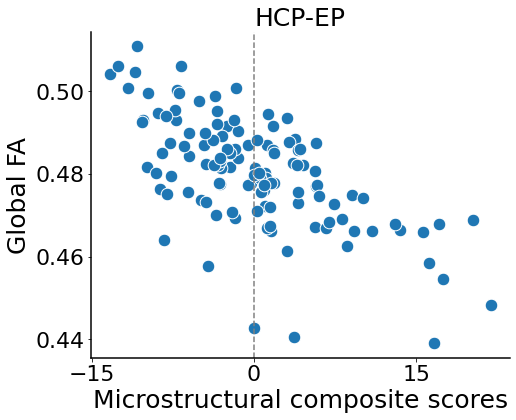
**A	B**

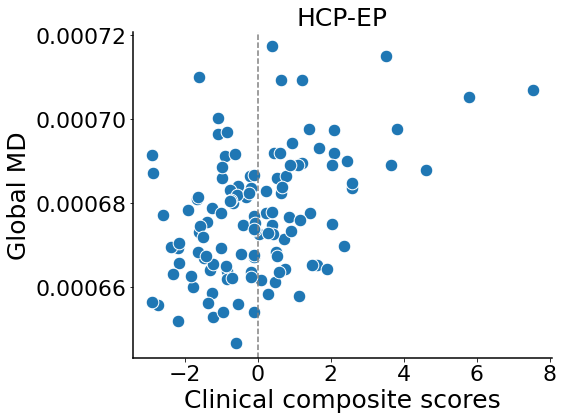

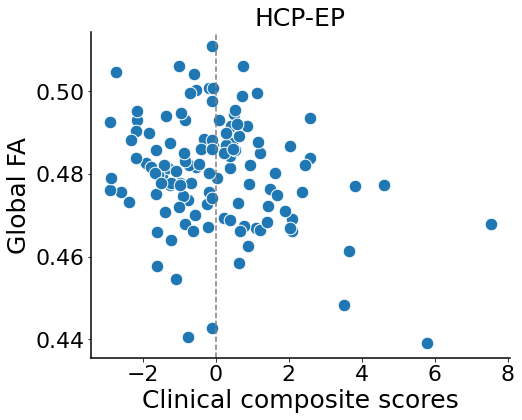
**C	D**

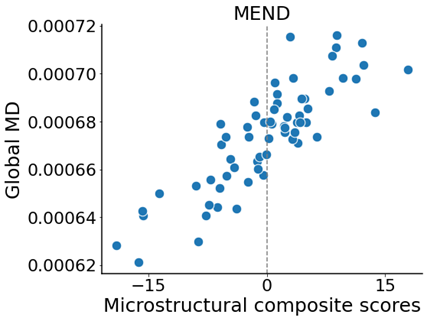

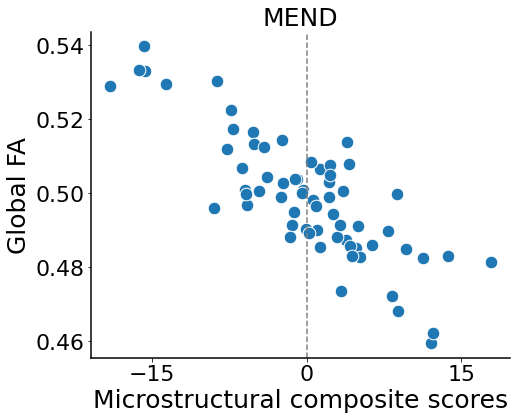
**E	F**

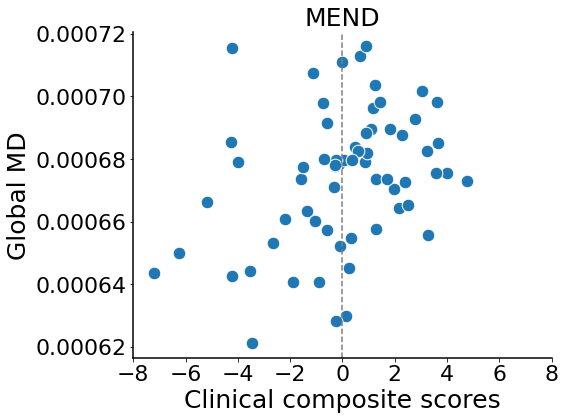

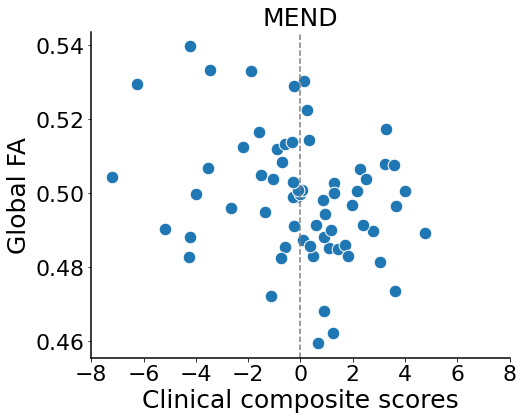
**G	H**

***Figure S4.*** *Loading coefficient by connectivity type of WM tracts.* Density and proportion of each tract category with respect to their loading coefficients in each cohort (Figure S4A-D). Density and proportion differences demonstrated through data exploration were primarily driven by the number of WM tracts in each connectivity category. No differing pattern of loading coefficients were observed by kernel density estimate.

**A	B**

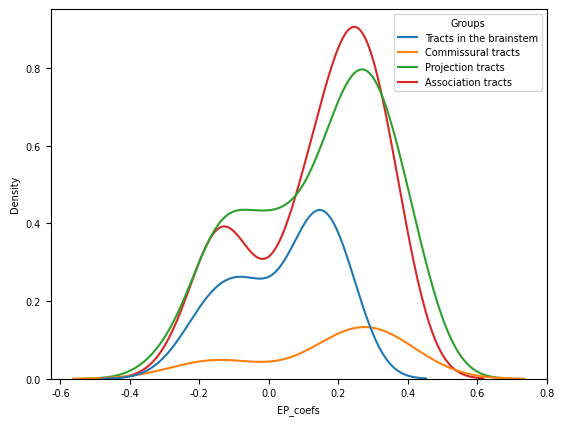

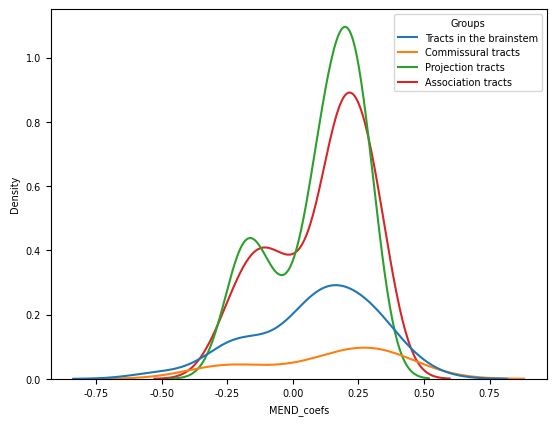

**C	D**

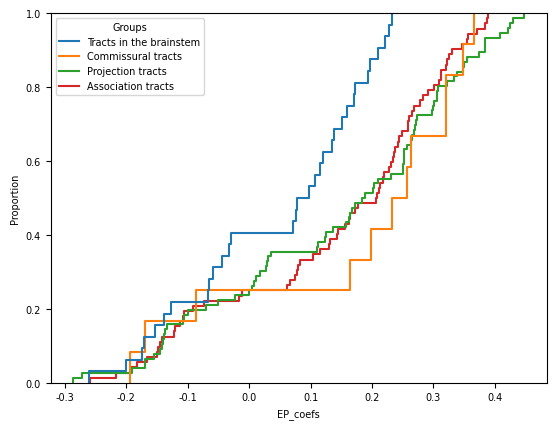

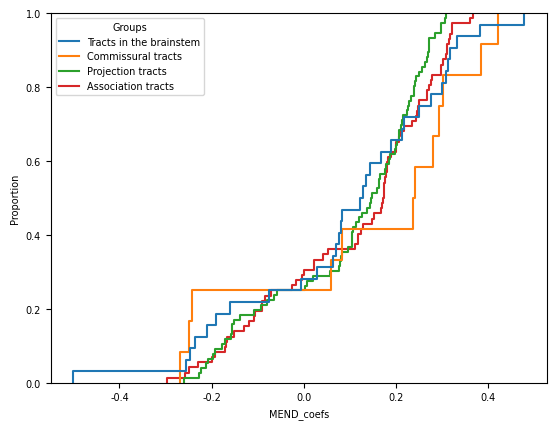

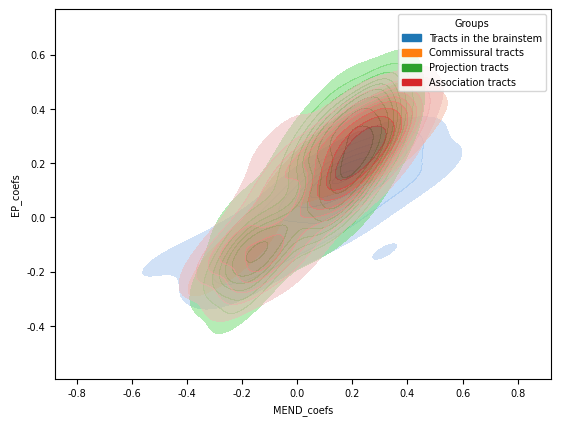
**F**
